# Extent of damage to descending output from cortex rather than to specific cortical regions drives the emergence of flexor synergy in non-human primates

**DOI:** 10.64898/2026.03.04.709517

**Authors:** Anna Baines, Isabel S. Glover, Anne M.E. Baker, John W. Krakauer, Stuart N. Baker

## Abstract

Obligate flexor synergies are a defining feature of the hemiparetic phenotype following stroke in humans. Although these intrusive synergies can diminish over time, recovery may plateau, leaving some individuals with movements permanently constrained to synergies. Despite their clinical significance, the neural mechanisms underlying the emergence and persistence of abnormal synergies remain poorly understood.

To investigate this mechanistically, three macaque monkeys were trained on a reach and grasp task prior to receiving one of three unilateral lesion types: 1) a focal sensorimotor cortical lesion, 2) a combined sensorimotor cortical and magnocellular red nucleus (RNm) lesion, or 3) a lesion of the internal capsule. Upper limb three-dimensional kinematics and EMG cross correlation were used to measure the intrusion of synergies during *in synergy* vs *out of synergy* reaching.

A combined RNm and cortical lesion produced weakness but no flexor synergy. A similar-sized cortical lesion generated mild synergies which substantially recovered. By contrast, a large internal capsule lesion produced severe, persistent flexor synergy. Collectively, these findings suggest that the emergence of abnormal synergies is determined by the extent of corticofugal disruption, and their persistence depends on the ability of surviving supraspinal motor pathways to regain selective control over muscle contractions.

## Introduction

A stroke affecting the middle cerebral artery causes damage to the sensorimotor cortex and its descending corticofugal projections, resulting in acute contralateral hemiparesis in approximately 80% of patients (Cramer et al., 1997; Hui et al., 2024). Motor recovery from such a stroke typically follows a stereotyped trajectory (Twitchell, 1951; Brunnstrom, 1970), beginning with weakness and loss of dexterity (negative signs) that evolve into spasticity and abnormal muscle synergies (positive signs). From this stage, patients either regain more coordinated, fractionated movement, or they remain constrained to synergistic movement patterns (Twitchell, 1951; Bourbonnais et al., 1989; Ellis et al., 2011).

The most prevalent abnormal pattern is the flexor synergy, in which voluntary shoulder abduction produces involuntary coactivation of elbow, wrist, and finger flexors, thereby limiting elbow extension and hand opening (Beer et al., 1999; Sukal et al., 2007; Ellis et al., 2008; Miller and Dewald, 2012; Ellis et al., 2017; Lan et al., 2017; Avni et al., 2024a). The Fugl-Meyer Assessment (FMA) was originally developed to quantify how constrained voluntary movements are by synergies, and remains one of the most widely used clinical outcome measures for upper limb impairment after stroke (Fugl et al., 1975).

To date, research has predominantly focused on restoration of strength and dexterity (negative signs) with less emphasis on identifying the neural circuits underlying abnormal synergies. Yet upper limb impairment persists in more than 50% of patients in the chronic phase of recovery (Coscia et al., 2019; Rodgers et al., 2019; O’Flaherty and Ali, 2024; Tang et al., 2024); synergies, not merely weakness or impaired dexterity, constitute a major barrier to functional recovery after stroke.

Following damage to the corticospinal tract (CST), recovery of upper limb function depends on reorganization of surviving motor pathways (Baker, 2011; Baker et al., 2015; Krakauer and Carmichael, 2017). When damage to the CST is substantial, the reticulospinal tract (RST) reorganizes its circuity in a compensatory attempt to restore lost input to spinal cord neurons (Baker, 2011; Zaaimi et al., 2012; Baker et al., 2015; Zaaimi et al., 2018; Karbasforoushan et al., 2019; Mooney et al., 2025). It seems, however, that this strategy prioritizes strength restoration at the expense of individual joint control and distal hand function (Sukal et al., 2007; Lan et al., 2017; Owen et al., 2017; McPherson et al., 2018b; Glover and Baker, 2020; Taga et al., 2024). Notably, increased reliance on the RST after stroke is associated with poorer measures of upper limb recovery (Stinear et al., 2007; Byblow et al., 2015; Choudhury et al., 2019; Schambra et al., 2019). Compared to the CST, the RST has more diffuse projections to the spinal cord (Kuypers et al., 1962; Matsuyama et al., 1997; Matsuyama et al., 1999; Lemon, 2008), which produce less fractionated upper limb movements (D G Lawrence, 1968a, b; Muir and Lemon, 1983; Baker and Perez, 2017; Tazoe and Perez, 2017; Glover and Baker, 2022) and demonstrate preferential facilitation of ipsilateral flexor muscles. It has therefore been suggested that the RST may be responsible for generating abnormal flexor synergy following stroke (Dewald et al., 1995; Beer et al., 2007; Ellis et al., 2007; Sukal et al., 2007; Lan et al., 2017; Owen et al., 2017; McPherson et al., 2018a).

One possibility is that the emergence of flexor synergies depends on which cortical regions are affected by the lesion. Specifically, synergies could result from damage to regions that would normally modulate reticulospinal output and prevent obligate, biased facilitation of the flexor synergy in healthy individuals. Past studies have highlighted the significance of an inhibitory cortical strip within the primary motor cortex (M1), termed ‘Area 4s’ (Hines, 1936; Hines, 1937, 1943), which projects to a portion of the reticular formation that is capable of exerting strong suppressive effects on motor output (Magoun and Rhines, 1946; McCulloch et al., 1946; Engberg et al., 1968; Jankowska et al., 1968; Takakusaki et al., 2001; Du Beau et al., 2012). Lesions to Area 4s in monkeys have produced strong positive signs in early reports from the 1940s (Hines, 1937, 1943; Denny-Brown and Botterell, 1948; Denny-Brown, 1966), but this was not replicated in our more recent study using lesions that produced varying degrees of damage to Area 4s in eight monkeys (Baines et al., 2026).

In the aforementioned study (Baines et al., 2026), lesions that produced varying degrees of damage to the premotor cortex (PMd), M1, and the primary somatosensory cortex (S1) also did not result in overt flexor synergies, even when combined with preceding unilateral lesions to the magnocellular red nucleus (RNm) in a subset of animals. These findings are consistent with clinical studies showing that cortical lesion size or location alone cannot reliably predict poorer FMA score (Shelton and Reding, 2001; Zhu et al., 2010; Page et al., 2013). Notably, the presence or absence of a lesion of the posterior limb of the internal capsule (PLIC) is the most reliable predictor of FMA improvement (Shelton and Reding, 2001), including the ability to recover isolated movements, which is rare after PLIC damage.

The PLIC contains densely packed axons from cortical motor and premotor areas that contribute to descending corticoreticular, corticorubral, corticopontine, and corticospinal tracts (Kuypers, 1960; Emos et al., 2023). Because of this dense convergence, even small capsular strokes have devastating motor consequences, and are unfortunately commonly encountered in clinical practice (Fries et al., 1993; Karbasforoushan et al., 2019). Following such damage to the PLIC and the ipsilesional corticospinal tract, patients rely on the contralesional cortex in attempts to restore some control over the paretic limb via the remaining motor pathways, chiefly the ipsilateral corticospinal tract and bilateral reticulospinal tract. Given that the ipsilateral corticospinal contributions to upper limb movements are minimal (Soteropoulos et al., 2011), and the RST inherently favors facilitation of upper limb flexors (Davidson and Buford, 2006; Herbert et al., 2010; Zaaimi et al., 2012), it is plausible that these surviving pathways are unable to suppress flexor synergy when ipsilesional corticospinal capacity is highly compromised (McPherson et al., 2018a). This could explain why increased contralesional cortical activity correlates with poorer upper limb recovery after stroke (Ward et al., 2003; Matsuura et al., 2017).

In this study, we set out to test if a lesion to the PLIC generates abnormal flexor synergy in a non-human primate model. First, we quantified changes in upper limb reaching performance using three-dimensional kinematic measures (Avni et al., 2024a) of flexion fraction proportion, flexion fraction strength, and path length. We compared performance of movements that required shoulder abduction paired with elbow extension (*out of synergy*) vs those that required shoulder abduction paired with elbow flexion (*in synergy*). Second, we assessed muscle coactivation strength with electromyography (EMG) cross correlation analyses. Third, we quantified the number of attempts required to initiate the different reach types.

All analysis methods were repeated using datasets from two monkeys which were included in our previous report (Baines et al., 2026). One animal had a large cortical lesion that spanned multiple sensorimotor regions (Monkey D); the second had a combined lesion of the RNm and multiple sensorimotor regions (Monkey Ca). We show that the internal capsule lesion (Monkey Cw) led to the poorest acute and chronic recovery outcomes with persistent flexor synergy. The pure cortical lesion generated weak flexor synergy expression, which resolved with further recovery. No evidence of the flexor synergy was observed following the combined RNm and cortical lesion.

## Materials and Methods

Experiments were conducted in one female (weight at start of study = 6.8 kg) and two male rhesus macaques (weights at start of study = 10.7 kg and 11.1 kg), referenced in this report as Monkey Ca, Cw and D, respectively. All animal procedures were carried out under an appropriate license issued by the UK Home Office under the Animals (Scientific Procedures) Act 1986 and were approved by the Animal Welfare and Ethical Review Board of Newcastle University.

### Behavioral Task

Animals were trained to perform a reach and grasp task with their right upper limb (Fig. 1A); this took between 3-6 months. The neck and left arm were gently restrained throughout. A trial of the task was initiated by the monkey holding a handle for one second, after which the lid of a baited cup rapidly retracted, providing access to a food reward contained within. The monkey then made a rapid reaching movement to retrieve the reward. The experimenter pressed a foot switch to signal the successful end of the trial. A timeout tone sounded if the footswitch was not pressed within 5s of handle release, and the trial would then be repeated from the start. Five or 10 repeat trials were performed in the same reach direction, after which the handle and cup were moved to another of the 12 possible arrangements on the vertices of a diamond.

**Figure 1.**
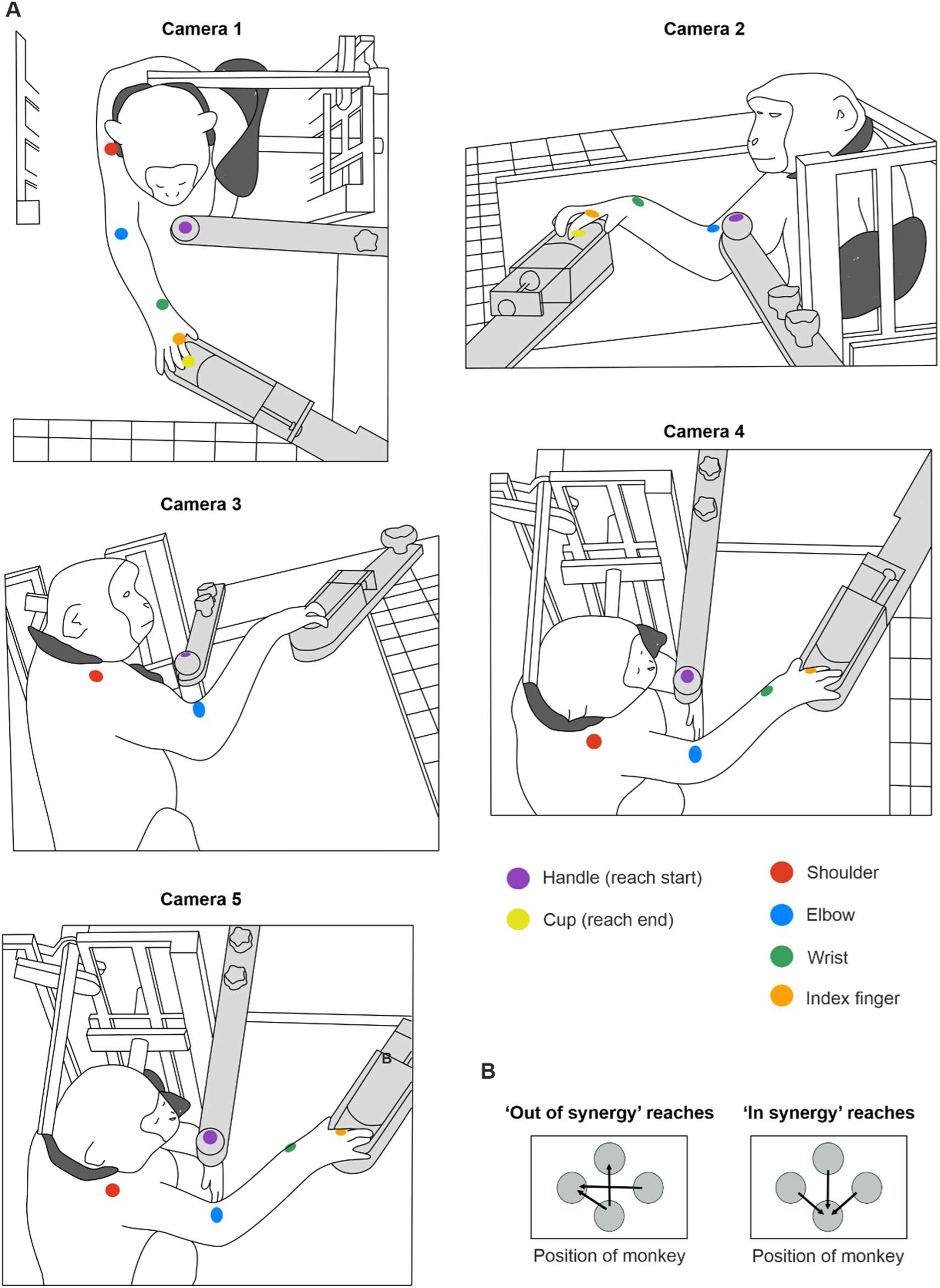
Behavioral task and 3D video recording configuration. A, drawings of the reaching task configuration from five different camera perspectives. Monkeys were placed in a neck and left arm restraint (dark grey). Handle and cup (light grey) positions were fixed to the table; here the trial has the handle near to the body and the cup further away, hence requiring shoulder abduction paired with elbow extension (an *out of synergy* reach). B, six possible handle and cup arrangements for assessing *out of synergy* versus *in synergy* reaches (determined as described in the text), shown from the perspective of camera 1 positioned directly above the monkey.

Each day, the animals performed an assortment of reaches requiring either shoulder abduction and elbow flexion (*in synergy* co-contractions) or shoulder abduction and elbow extension (*out of synergy* co-contractions). Figure 1A demonstrates an *out of synergy* reaching movement, which would be extremely difficult to perform with abnormal flexor synergy expression.

### Surgical Preparation

Once animals were fully trained, they underwent an aseptic surgery to implant chronic EMG electrodes and a headpiece.

All surgeries described in this report were conducted under general anesthesia managed by a veterinary surgeon. Monkeys were initially sedated with an intramuscular injection of ketamine (2-10 mg/kg), medetomidine (3 μg/kg), and midazolam (0.3 mg/kg). During surgical preparation, anesthesia was maintained with sevoflurane inhalation (2-3%). Animals were intubated and ventilated with positive pressure ventilation. Where electrophysiological recordings of single units in the cortex were made to map the morphology of the central sulcus, anesthesia was switched to a continuous infusion of alfentanil (24-108 µg/kg/hr), ketamine (6 mg/kg/hr), midazolam (0.3 mg/kg/hr), as we have found that this better preserves cortical activity. Infusions of methylprednisolone (5.4 mg/kg/hr) and Hartmann’s solution (5 ml/kg/hr) were used to prevent edema and dehydration respectively. The animal was kept warm with a temperature-controlled heating blanket, and a supply of thermostatically-controlled warm air. Vital signs were monitored, including pulse oximetry, heart rate, blood pressure, core and peripheral blood temperature, and end tidal CO_2_. Anesthetic doses were adjusted as necessary to maintain a stable plane of deep anesthesia.

Pairs of Teflon-insulated stainless-steel wires (AS632, Cooner Wire Co, Chatsworth, CA, USA) were implanted into right upper limb muscles including the anterior deltoid, posterior deltoid, supraspinatus, triceps brachii, biceps brachii, brachialis, and brachioradialis. EMG wires ran subcutaneously to connectors (A22004-001, Genalog, Kent, UK) that were cemented to a headpiece. To construct the headpiece, ceramic screws were fitted in the skull to support three plastic posts. Dental acrylic was built up on top of the skull, and around the skull screws, posts, and EMG connectors. A suitable recovery period was allowed to promote wound healing (usually two weeks), and post-operative therapy given as appropriate including broad spectrum antibiotics (oral or subcutaneous co-amoxyclav 12.5 mg/kg), analgesics (intramuscular buprenorphine 0.02 mg/kg; oral or intramuscular meloxicam 0.2 mg/kg; oral paracetamol 23 mg/kg), and steroids (intramuscular dexamethasone 0.2-0.5 mg/kg).

A second surgery was performed two weeks later in Monkey Cw. A parylene-insulated tungsten electrode (LF501G, Microprobe) was implanted chronically into the left pyramidal tract (PT), rostral to the pyramid decussation. The electrode was passed through a small convenient craniotomy made during surgery, and a double-angle stereotaxic technique applied to aim the electrode at the target, with reference to the craniotomy as the starting point (Soteropoulos and Baker, 2006). Optimal fixing locations were determined with reference to antidromic cortical responses and spinal volleys elicited by stimulation through the electrode (500µA 0.1ms biphasic pulses); volleys were recorded from epidural wires placed over left and right M1, and from a wire temporarily inserted using a needle near to the cervical spine. The optimal fixing location was determined as the lowest threshold required to elicit an antidromic response in ipsilateral M1, with no response from contralateral M1. The electrode was then connected to a connector, and dental acrylic used to fix both electrode and connector to the headpiece.

Appropriate post-operative care was given, and the animal was well enough to return to the laboratory two days later for baseline recording. The electrode implant enabled selective stimulation of the left PT during recording sessions to assess motor evoked potentials in implanted upper limb muscles.

### Three-Dimensional Markerless Joint Tracking

Baseline data were collected five days per week, for three to five weeks, to sample normal reaching behavior before lesions were made. Further kinematic data were then gathered for 15 weeks.

High-speed video footage was recorded simultaneously using five cameras positioned around the monkey (90–100 fps), as illustrated in Fig. 1A. Three cameras recorded at a resolution of 1280 × 1024 pixels (CM3-U3-31S4C-CS, 1/1.8” Chameleon®3 Color Camera, Teledyne FLIR, Oregon, USA), and two cameras recorded at 1920 × 1200 pixels (Blackfly S BFS-U3-23S3C, Teledyne FLIR, Oregon, USA). Images were acquired with either FlyCap or SpinView GUI (Spinnaker SDK software, Teledyne FLIR, Oregon, USA). Camera positions were chosen to guarantee consistent visibility of the targeted landmarks from a minimum of two cameras throughout the session. Simultaneous frame capture was triggered by regular digital pulses sent to each camera from a custom circuit containing an Arduino microcontroller. The microcontroller also generated an 8-bit binary counter which incremented once with each trigger; the counter output was displayed via eight LEDs visible within the camera frame. On occasion, the camera software missed writing a frame to disk. We identified such missed frames using a custom MATLAB script which extracted and thresholded the light intensities around each LED to determine the binary count and detect non-contiguous frames.

After each session, videos of a moving 9 x 6 checkerboard were recorded simultaneously from all five cameras to calibrate camera positions in the later analysis pipeline. Task videos, calibration videos, and task parameters were stored to disk.

We used the open source markerless tracking software DeepLabCut (v2.2, (Nath et al., 2019)) to track two-dimensional locations of the handle (start of reach), cup (end of reach), elbow (acromioclavicular joint, a defined location of the skin at the inner elbow), wrist (ulnar styloid process), and index finger (the right index finger metacarpophalangeal joint) (Fig. 1A). Video frames required to train the deep neural networks were extracted using a k-means algorithm from all five cameras, and landmark positions manually labelled in each extracted frame. Landmark positions were marked with animal identification pen ink (Monkey Ca) or permanent tattoos (Monkeys D and Cw) to improve manual labelling accuracy and consistency across sessions. Separate tracking models were used for Monkey Ca vs. Monkey D and Cw due to different markings (ink vs tattoos).

Two Res-Net 50 models, pre-trained on ImageNet, were custom-trained on manually-labelled frames from all five camera perspectives (Monkey Ca – 2299 frames; Monkeys Cw and D – 1302 frames). Using a prediction likelihood cutoff of 0.6, the trained networks achieved a training error of 2.17 mm and a test error of 2.98 mm for Monkey Ca, and a training error of 2.25 mm and a test error of 2.98 mm for Monkeys D and Cw. All session videos were processed offline using the appropriate trained DLC network.

For all monkeys, in some cases, we detected non-contiguous frame indices in task and calibration videos resulting from a failure of the software to keep up with writing the data stream to disk; these needed to be accounted for prior to 3D joint reconstruction. For calibration videos, new videos were generated in MATLAB using only frames captured simultaneously by all five cameras. For task videos with non-contiguous frames, the arrays containing tracking predictions for each video frame output by DeepLabCut were corrected using a custom Python script. Placeholder rows were automatically inserted at the missing frame indices containing interpolated 2D predictions and prediction likelihood values set to zero (meaning these frames would be ignored in later processing). This process ensured all arrays (one per camera) had consistent frame counts.

Aligned calibration videos and aligned files containing DeepLabCut predictions were used to reconstruct three-dimensional trajectories of the shoulder, elbow, and wrist joints for each session using Anipose (v1.0.1; (Karashchuk et al., 2021)), an open-source Python toolkit that builds on 2D DeepLabCut outputs. The five cameras were calibrated by detecting checkerboard key points and estimating intrinsic and extrinsic camera parameters with iterative bundle adjustment (Karashchuk et al., 2021). Anipose used the calibrated parameters to triangulate the multi-view 2D DeepLabCut predictions and obtain 3D anatomical landmark positions. This process was followed by temporal optimization to reduce frame-to-frame noise.

### Kinematic Analysis

All analyses were performed off-line using custom software written in MATLAB. The start of each trial was defined by the release of the handle contact. The monkey then made reaching movement towards the cup to retrieve a piece of food. Pre-lesion, the animals reliably made a single successful reaching movement. However, post-lesion, performance could be significantly impaired and it could take multiple cyclical reaching movements for the animals to reach the cup. Therefore, to provide a comparable metric between pre-lesion and post-lesion reaches, we focused our analysis on the first reaching movement made towards the cup (the ‘initial reach’). The start of the reach was defined by release of the handle, and the end of the reach defined as the first transition from deceleration to acceleration (zero velocity) after the halfway point between handle and cup was exceeded. This analysis was undertaken using 2D measures of index finger position output by DeepLabCut using the top camera view (Fig. 2A).

**Figure 2.**
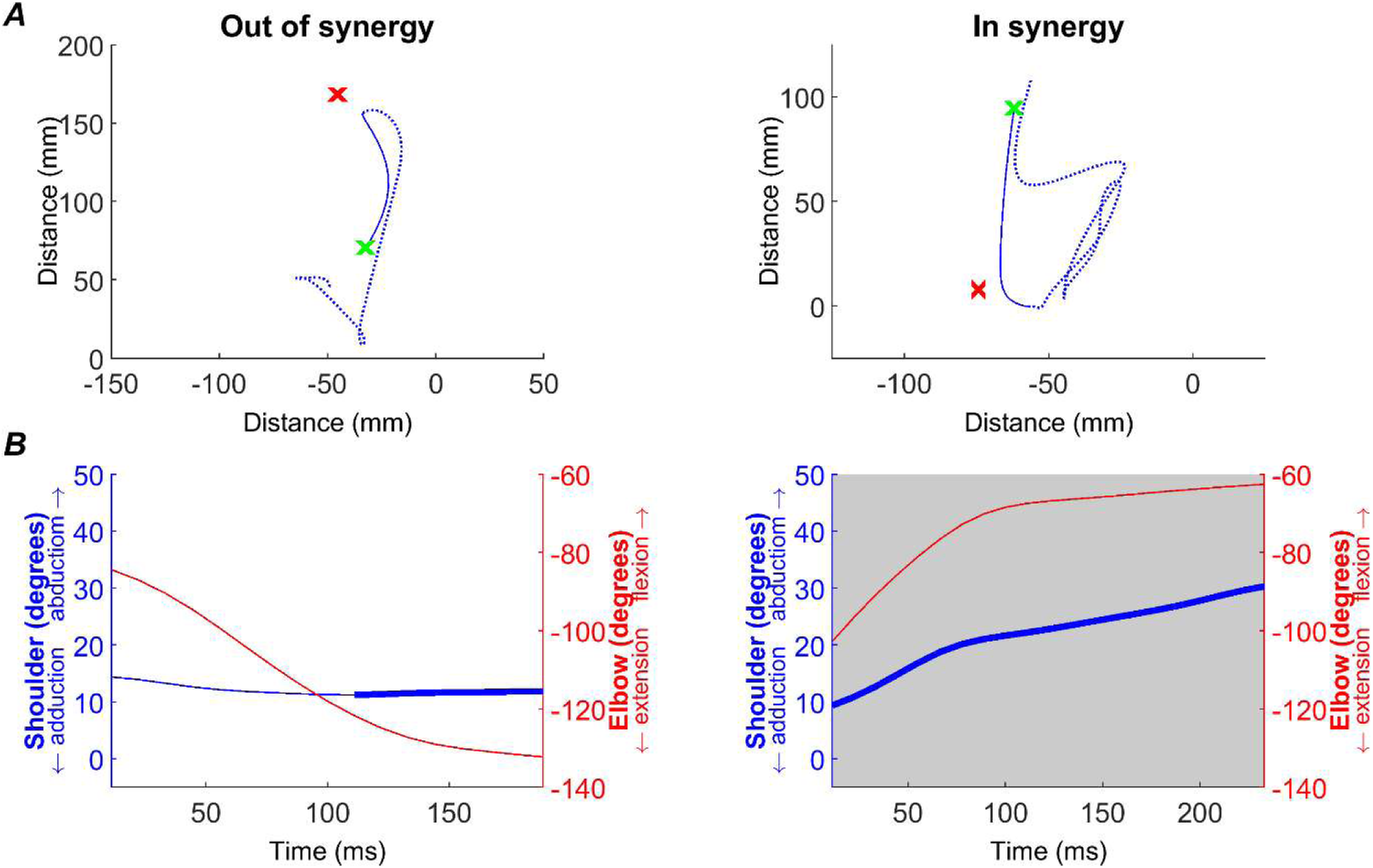
Kinematics measures. Example of kinematic analysis, demonstrated with a single *out of synergy* (first column) and *in synergy* (second column) trial. A, path taken from handle (green x) to cup (red x), as measured by index finger tracking. The full movement recorded for the trial is shown by the dotted blue line, whilst the solid line shows the initial reach (see Methods) that was used in all subsequent analyses. B, flexion-fraction proportion and strength were calculated by measuring shoulder abduction/adduction (blue trace, left axis) and elbow flexion/extension (red trace, right axis) during the initial reach. During the longest period of shoulder abduction (thick blue trace), the correlation between shoulder and elbow angles was calculated to give flexion-fraction strength (see Methods). For the shown traces, this was -0.98 for the *out of synergy* movement and 0.99 for the *in synergy* movement. Flexion-fraction proportion was calculated as the proportion of the initial reach with shoulder abduction and elbow flexion (grey shaded region). For the traces shown, flexion-fraction proportion was 0 and 1 for the *out of synergy* and *in synergy* reaches, respectively. Data from monkey Cw baseline period.

To investigate the relationship between movements around the shoulder and elbow joints, we used 3D coordinates to construct joint angles. If **r_s_**, **r_e_** and **r_w_** are vectors representing the detected 3D coordinates of shoulder, elbow and wrist markers respectively, then the angle between upper arm and forearm θ, which is the angle of elbow flexion, was calculated as:

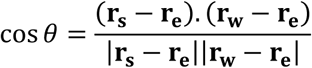

Where . represents the vector dot product, and |•| the vector magnitude. For the shoulder, we formed a vector **r_se_** representing the component of upper arm movement in the adduction-abduction direction:

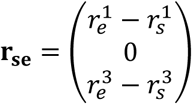

Where the three canonical axis directions were (in order) in the transverse, sagittal and longitudinal directions. Shoulder abduction angle φ was calculated as:

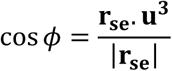

Where **u^3^**is the unit vector in the vertical direction:

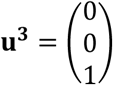

We then replicated the two metrics reported by Avni et al. (2024b): flexion-fraction proportion and flexion-fraction strength. We performed this analysis using elbow flexion/extension and shoulder abduction/adduction as these movements were reliably activated during our task and reflect the presentation of synergies in stroke survivors.

Flexion-fraction proportion was calculated as the proportion of the movement time in which both the elbow was flexing and the shoulder abducting. This was measured across the initial reach period, as defined above (Fig. 2B; grey region). Flexion-fraction strength was the correlation between shoulder and elbow angles during the longest continuous period of shoulder abduction (Fig. 2B, bold blue trace). Since the monkeys often took indirect paths to the cup post-lesion, we limited this analysis to the greatest amplitude shoulder abduction in which movement was made in the direction towards the cup (within ±90° of handle to cup direction). Continuous elbow flexion during this period of shoulder abduction would yield a flexion-fraction strength of +1 (an *in synergy* co-contraction), whilst continuous elbow extension during shoulder abduction would yield a flexion-fraction strength of -1 (*out of synergy* co-contraction).

Finally, to assess overall performance on the task, we also measured the path distance as the total distance travelled by the monkey’s index finger for the initial reach.

All combinations of the four possible positions for cup and handle produced 12 different reaches that the monkeys were trained to complete. To provide a comparison to the human literature (Avni et al., 2024b), we restricted our analysis to *in synergy* and *out of synergy* reaches. To identify these, prior to any lesions we analyzed all 12 reaching directions, calculated the flexion-fraction strength and then selected the reaching tasks with flexion-fraction strengths approaching +1 for *in synergy* movements and approaching -1 for *out of synergy* movements. This approach identified three *in synergy* and three *out of synergy* reaching directions (Fig. 1B). We combined these movements in subsequent analysis to provide *in synergy* and *out of synergy* metrics.

In Monkey Cw, we observed that following the lesion surgery, the animal not only struggled to perform the reaching movements required of the task, but also struggled to hold onto the handle that initiated each trial. Although the monkey was able to touch the handle briefly, he struggled to maintain contact with the handle for the required hold period. To quantify this impairment, for each session we counted the number of ‘hold attempts’ prior to a successful 1 s-long hold period that opened the food cup. Since the subsequent reaching task was irrelevant here, we focused this analysis on all trials with the handle in the positions nearest to and furthest away from the animal.

All recording sessions were grouped by week relative to the lesion, with the first post-lesion week being week zero. Baseline/pre-lesion parameters were calculated by averaging all sessions prior to the first lesion received by each monkey. Each post-lesion time point was compared to the pre-lesion average with unpaired t-tests, and multiple comparisons corrected using the Benjamini-Hochberg procedure with a false discovery rate of 5%. These analyses were repeated grouping the recording sessions into pre-lesion, subacute (0-5 weeks post-lesion) and chronic (10-15 weeks post-lesion) phases. For both weekly and phase analysis, averages were only calculated if there were data from at least five trials.

### Motor Evoked Potentials following Left Pyramidal Tract Stimulation in Monkey Cw

In Monkey Cw, motor evoked potentials (MEPs) following left PT stimulation were recorded from the right posterior deltoid and brachioradialis muscles while the animal was freely reaching for food. Each recording session consisted of 200 repeated PT stimuli at 2 Hz frequency, stimulus intensity 700 μA. Baseline data were collected five days per week for five weeks to sample normal reaching behavior before the lesion. Subsequent recordings were collected for 15 weeks post-lesion to assess changes in MEP size over time.

Amplified EMG signals sampled at 5 kHz were processed offline using custom-written MATLAB scripts (MathWorks, Massachusetts, US). Raw EMG was full wave rectified and smoothed (σ = 0.5 ms). For each muscle, and each recording session, EMG traces from 50 ms before to 50 ms after each PT stimulus were overlaid and visually inspected. Traces containing large electrical artefacts were excluded from further analysis. The remaining sweeps were averaged for each recording session and overlaid across all sessions to allow accurate manual placement of two cursors defining the onset and offset latency of the MEP. The area under the curve (AUC) was then calculated for each session and averaged per week.

### EMG Cross Correlation

This analysis quantified the strength of muscle coactivation patterns (synergies) that were the opposite to those required to support the intended movement, and whether synergy expression changed post-lesion. For *out of synergy* movements which required elbow extension, we calculated the coefficient of determination (r^2^) between shoulder abductors and elbow flexors (i.e. the flexor synergy) across all monkeys. For *in synergy* movements which required elbow flexion, the same analysis was performed between shoulder abductors and an elbow extensor (triceps); however, this analysis was only possible for Monkey Cw due to technical issues with the triceps EMG recordings in the other two animals.

EMG for each muscle and recording session was initially inspected visually, and muscles were only included if recorded data were consistently free of electrical artefacts and had minimal, consistent-amplitude noise. EMG recordings were available from the following muscles: monkey Cw, anterior deltoid, supraspinatus, biceps, brachioradialis, brachialis, and triceps; monkey D, posterior deltoid, biceps, brachioradialis, and brachialis; monkey Ca, anterior and posterior deltoids, biceps, brachioradialis, and brachialis.

EMG data were processed offline using custom-written MATLAB scripts (MathWorks, Massachusetts, US). EMG recorded during each trial was extracted from a window spanning 0.2s before to 0.5 s after the time of handle release, then full wave rectified and smoothed using a Gaussian kernel (width 20 ms). Trial data for each muscle were only used if the maximum peak-to-peak EMG was smaller than the mean EMG value plus five standard deviations; this excluded electrical noise artifacts.

Cross correlation coefficients were computed between each possible shoulder abductor and elbow flexor or extensor EMG pairing. Values for each baseline and post-lesion week were averaged over all muscle pairs. The difference in r^2^ for each post-lesion week compared to baseline average r^2^ was found. A Monte Carlo approach then calculated the significance of these differences. All data from baseline and each post-lesion week for each muscle pair in turn were merged, shuffled, and reassigned to either a simulated pre-lesion or post-lesion dataset, and the r^2^ difference calculated. This process was repeated with 1000 iterations for each post-lesion week and each muscle pair to achieve a distribution of expected r^2^ differences under the null hypothesis of no difference between baseline and post-lesion. The actual difference in r^2^ for this post-lesion week vs baseline was compared to this distribution of shuffled e^2^ values, and was deemed statistically significant if p<0.05.

### Internal Capsule Lesion

In Monkey Cw, a unilateral lesion of the left internal capsule was performed using thermocoagulation under general anesthesia.

A craniotomy was made stretching from 16-25 mm anterior to interaural line (IAL) and 6-13 mm to the left of the midline. Burr holes were drilled over left M1 (19 mm anterior to IAL, 18 mm lateral) and SMA (25 mm anterior, 2mm lateral) to record antidromic volleys with epidural ball electrodes.

We began by mapping the left internal capsule with a parylene-insulated tungsten (LF501G, MicroProbe, San Jose, USA), targeting initial coordinates estimated from a structural MRI conducted prior to the surgery. A series of penetrations at various anterior-posterior locations were made, and single pulse stimuli (biphasic, 0.1 ms per phase) were delivered at each site. When antidromic volleys were recorded from left M1, the stimulation intensity was reduced to determine the threshold. Stereotaxic coordinates and corresponding stimulation thresholds were noted on graph paper, producing a fine-grained map of the internal capsule. Nineteen lesion locations were chosen, determined as the lowest threshold sites. These were located between 16-20 mm anterior, 6.6-8.6 mm lateral, and 13.0-20.0 mm dorsal to IAL.

Before making any lesions, baseline antidromic volleys were recorded from M1 and SMA following stimulation through the chronically implanted PT electrode (single shock, 1 mA, 0.07 ms pulse width). A neuromuscular blocker (atracurium 0.75 mg/kg/hr IV) was given to prevent muscle twitches during lesioning, which could potentially cause unwanted vibration of the electrode. Arterial blood pressure was monitored continually via a catheter inserted into the femoral artery; this allowed us to ensure anesthesia was stable even though muscle contractions were blocked. Lesions were then made using a thermocoagulation probe (TC-0.7-2-200, Cosman Medical Inc) connected to a radiofrequency generator (Radionics RFG-5). At each chosen site radiofrequency current was passed to coagulate tissue around the tip. After each lesion, antidromic volleys in left M1 and SMA were checked again. We continued producing lesions until the antidromic volleys were nearly completely abolished. The lesion and recording electrodes were then removed, craniotomies sealed, and the arterial line removed.

The animal was moved to a warm, padded recovery cage. Appropriate post-operative therapy was given, including broad spectrum antibiotics (oral or subcutaneous co-amoxyclav 12.5 mg/kg), analgesics (intramuscular buprenorphine 0.02 mg/kg; oral or intramuscular meloxicam 0.2 mg/kg; oral paracetamol 23 mg/kg) and steroids (intramuscular dexamethasone 0.2-0.5 mg/ml). In this animal, intravenous and subcutaneous fluid interventions were necessary for the first 7 days post-lesion due to facial weakness that restricted natural fluid consumption.

Recovery in Monkey Cw was markedly more protracted than in Monkey Ca and D, as was anticipated given the complete nature of the lesion. Symptoms included hemiparesis affecting the entire left side of the body (face, trunk, upper and lower limbs), as well as apathy, fatigue, and loss of appetite. Laboratory recording restarted 9 days post-lesion, once the animal was able to chew and swallow, was sufficiently hydrated, could safely move into the training cage, and was motivated enough to work despite the disability.

### RNm Lesion

A unilateral electrocoagulation lesion of the left RNm was conducted in Monkey Ca. The surgical method is described in detail in our earlier report (Baines et al., 2026).

### Cortical Lesions

Lesions using controlled injections of endothelin-1 were performed in Monkey Ca and D. This caused potent vasoconstriction in surrounding tissue, thus mimicking the effects of an ischemic stroke. Detailed surgical methods for these cortical lesions are available in our earlier report (Baines et al., 2026); only brief details will be given here.

In Monkey Ca, we performed a large cortical lesion 7 weeks after the RNm lesion. Endothelin-1 was infused 1-7 mm anterior to the central sulcus, with the intention of damaging the entirety of ‘Old M1’ (Rathelot and Strick, 2009). In Monkey D, we performed a more extensive cortical lesion which targeted the dorsal premotor area (PMd), M1 and the primary somatosensory cortex (S1; Brodmann’s Areas 1, 2 and 3). In both animals, injections covered the upper limb representation (8-19 mm lateral from midline). Appropriate post-operative care was given following lesions surgeries. Both animals were well enough to return to the laboratory for recording two days post-lesion.

### Histology

At 15 weeks post-lesion animals were terminally anesthetized with a lethal dose of propofol and perfused through the heart with phosphate buffered saline, followed by formalin for fixation of tissues. The brain and brainstem were removed and immersed in formalin for 24 hours before being transferred through ascending concentrations of sucrose solution (10%, 20%, 30%) for cryoprotection.

In Monkey Cw, the brain containing the left and right internal capsule was dissected into blocks and transverse sections (50 µm thick) were cut on a cryostat. Sections were stained using cresyl violet or nuclear fast red and Luxol fast blue to highlight grey and white matter respectively. Brightfield images of all stained sections were visually inspected using OMERO Plus Software (v5.29.2, Allan et al., 2012), and anatomical landmarks were compared against an interactive macaque Brain Atlas (Mikula et al., 2007). Outlines of sections spanning the complete dorso-ventral extent of the lesion were produced. In Monkey Ca, coronal midbrain sections were stained with cresyl violet and Perl’s Prussian blue with nuclear fast red to verify the locations of the three electrolytic lesions in RNm. In both Monkey Ca and D, post-mortem histological analysis was conducted to confirm the extent of the ischemic lesions. Parasagittal sections from the left and right sensorimotor cortices were stained with cresyl violet, and cytomorphology was examined in lesioned (left) versus non-lesioned (right) tissue. Full histological methods have been previously described in our earlier report (Baines et al., 2026).

## Results

### Kinematic Assessments of Reaching

This study aimed to compare metrics of pathological synergies after three different types of lesions: damage to the sensorimotor cortex alone, cortical damage combined with a preceding RNm lesion, or a lesion to the internal capsule.

Figure 1 shows the five-camera configuration we used to collect high speed video footage of the animals performing the reaching task. In this example, the monkey is performing an *out of synergy* movement that required shoulder abduction combined with elbow *extension*. By contrast, an *in synergy* movement involves shoulder abduction combined with elbow *flexion*, returning the arm towards the body. The positions of landmarks tracked in 3D using Deeplabcut (Nath et al., 2019) and Anipose (Karashchuk et al., 2021) pipelines are indicated by colored circles. Landmarks included the handle, cup, acromioclavicular joint, inner elbow, ulnar styloid process, and the metacarpophalangeal joint of the right index finger. This tracking approach enabled extraction of shoulder abduction-adduction, and elbow flexion-extension changes across different reaching movements, both in a healthy baseline period and for 15 weeks following each lesion.

Figure 1B presents the handle and cup arrangements, shown from the perspective of camera 1 placed directly above the monkey. The handle represents the start of the reaching movement, and the cup the end. We analyzed the performance of three *in synergy* reaches, which would require elbow flexion, and three ‘out of synergy’ reaches which would require elbow extension.

### Extent of Internal Capsule Lesion

Our aim was to lesion the posterior limb of the left internal capsule selectively using thermocoagulation, as this region contains descending corticospinal fibers from M1, as well as ascending, sensory thalamocortical fibers (Ara and Islam, 2010; Emos et al., 2023). During the lesion surgery, we monitored antidromic volleys following pyramidal tract stimulation as a way of assessing the damage to the corticospinal tract (Fig. 3A). After successive internal capsule lesions, the antidromic volley recorded from both M1 and SMA was markedly reduced (red traces, Fig. 3B).

**Figure 3.**
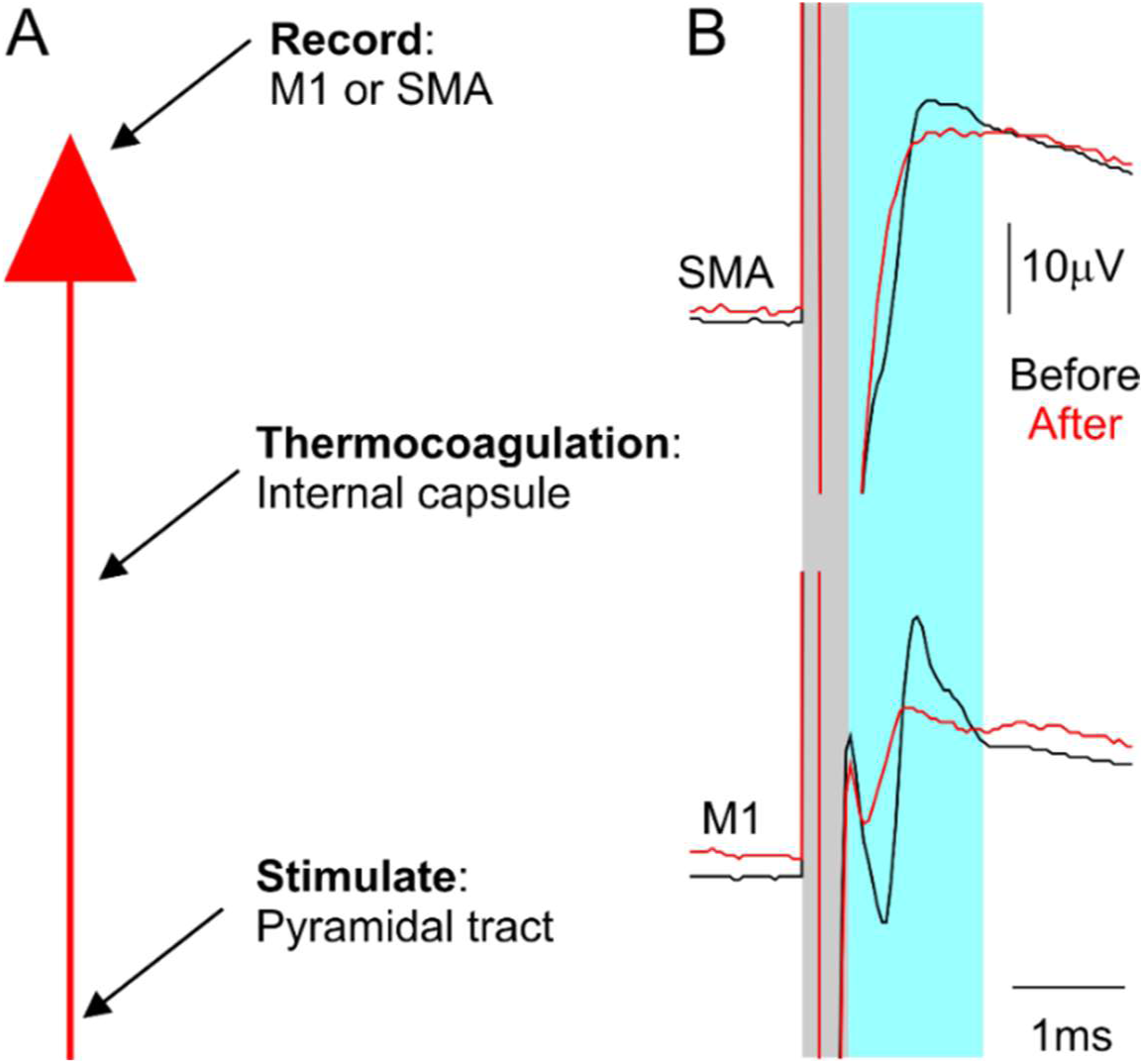
Electrophysiological recording used to guide targeted lesioning of the posterior limb of the left internal capsule. A, diagram showing the electrophysiological recording process used to perform and check internal capsule lesion efficacy. B, example electrophysiological traces from SMA (top) and M1 (bottom), recorded before (before) and after (red) the lesions. Grey regions represent the stimulus artefact; blue regions represent the elicited antidromic cortical volleys.

Figure 4 shows motor evoked potentials (MEPs) recorded from the right brachioradialis and posterior deltoid muscles following stimulation of the PT in the conscious state in Monkey Cw. The area under the curve of MEPs evoked by each stimulus was calculated and averaged per week of recordings (Fig. 4A). Smoothed and rectified EMG traces averaged over 200 stimuli from individual sessions during the baseline (blue), subacute (red) and chronic (pink) recovery periods are also shown (Fig. 4B). Following the internal capsule lesion, MEPs from both muscles were reduced to near zero and did not recover (pink and red traces) This indicates that the lesion caused permanent disruption of corticospinal tract connectivity to these *in synergy* upper limb muscles.

**Figure 4.**
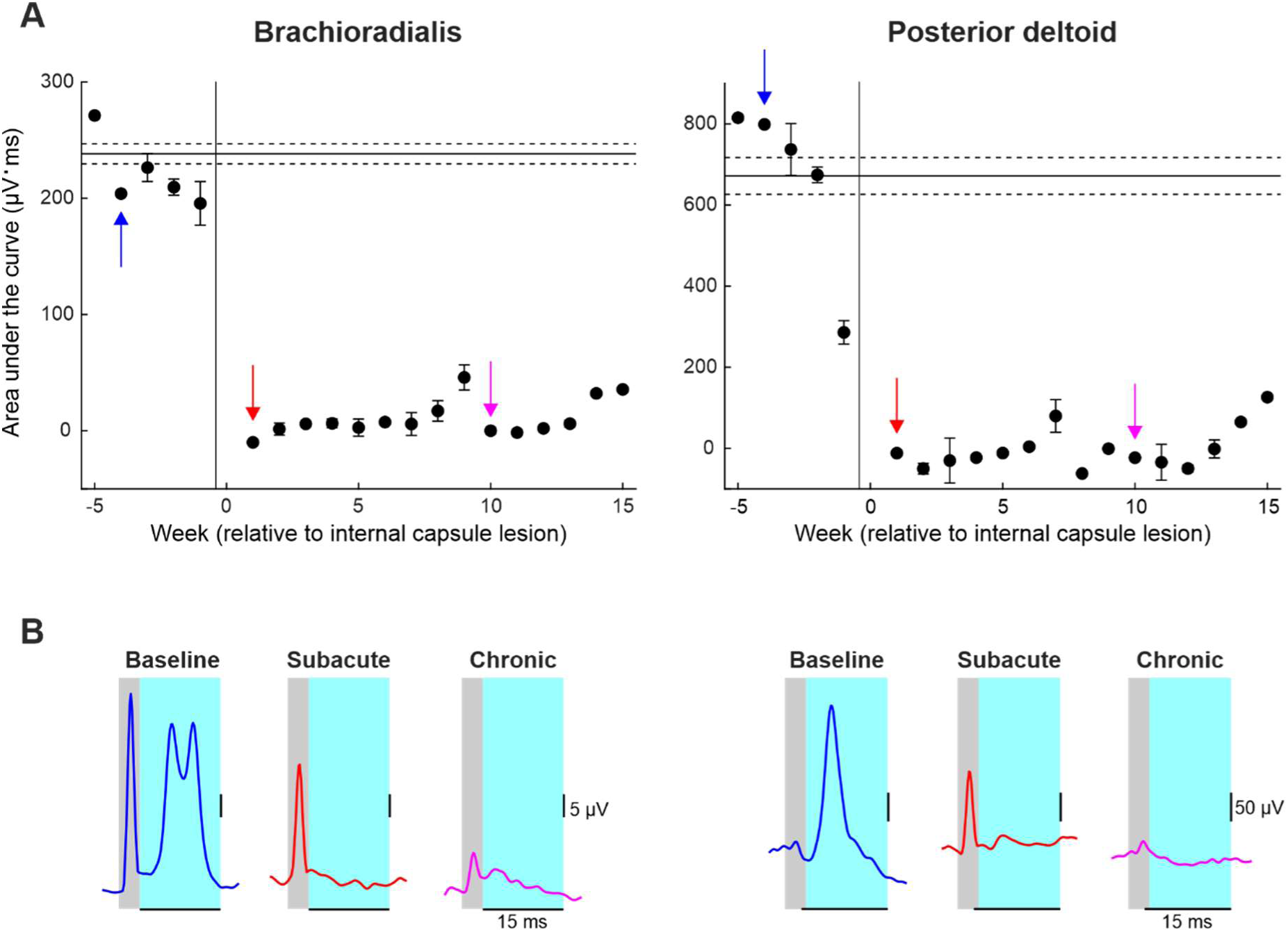
Motor evoked potential (MEP) recordings from the right brachioradialis and posterior deltoid following left pyramidal tract stimulation in Monkey Cw. A, area under the curve (AUC) of MEPs recorded from the brachioradialis (left) and posterior deltoid (right), shown as weekly averages. The vertical line indicates the time of the internal capsule lesion; horizontal lines indicate baseline mean ± standard error. Colored arrows indicate weeks corresponding to example traces shown in B. B, smoothed, rectified EMG traces averaged over 200 stimuli from individual sessions during the baseline (blue), subacute (red) and chronic (pink) recovery periods. Grey regions represent the stimulus artefact; cyan regions represent the time window over which the MEP AUC was calculated.

Figure 5 presents post-mortem histological analysis of horizontal brain sections used to confirm the extent of the internal capsule lesion in Monkey Cw. Figure 5A is an example left hemispheric section containing the internal capsule and surrounding structures, stained with luxol fast blue and cresyl violet for visualization of white and grey matter respectively. The lesion cavity can be seen in the center of the section with an approximate cross-sectional area of 4.5 mm^2^.

**Figure 5.**
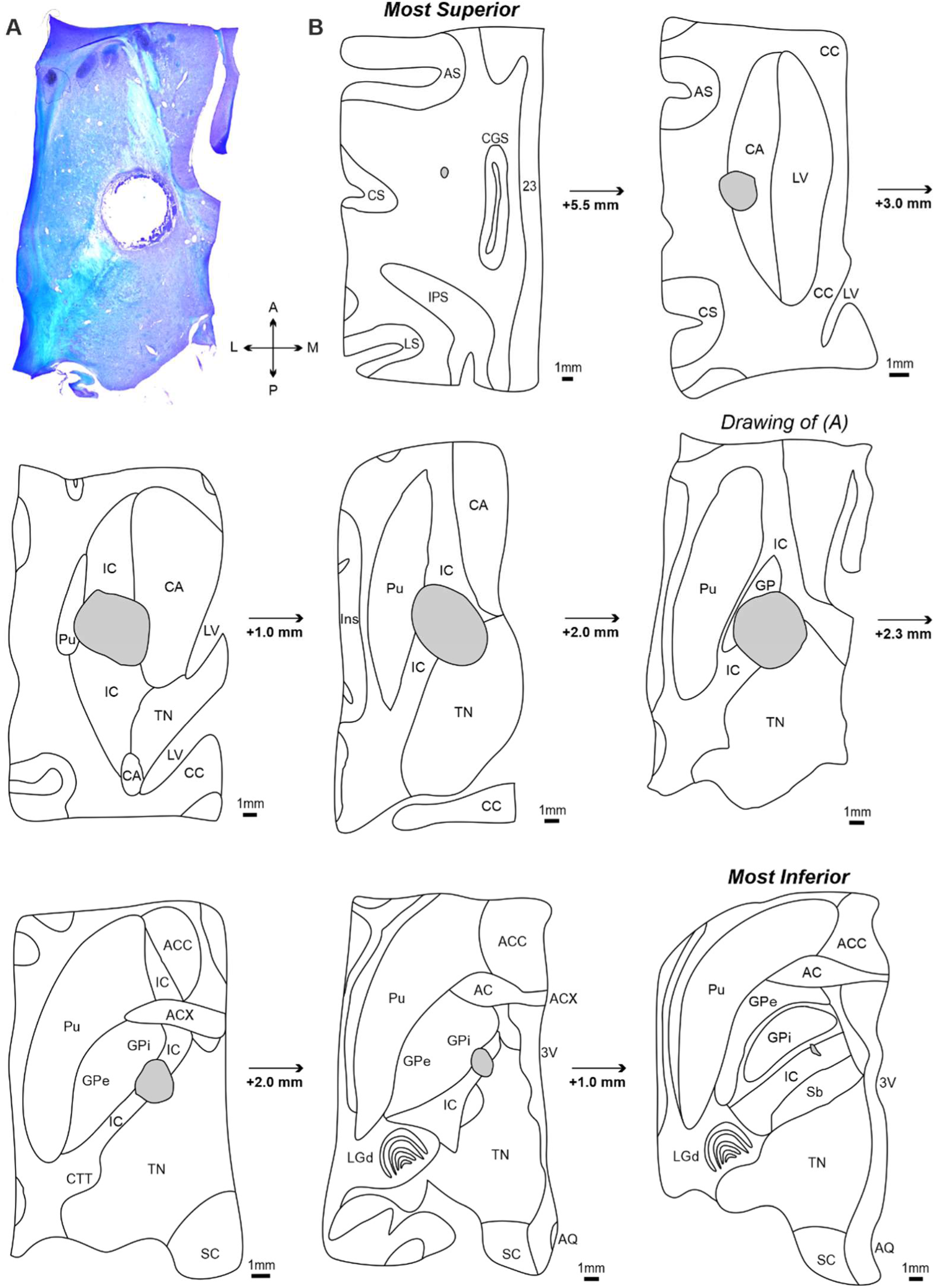
Confirmation of internal capsule lesion extent in Monkey Cw. A, transverse section from the left hemisphere containing the internal capsule and basal ganglia, counterstained with cresyl violet and luxol fast blue for comparison of grey vs white matter, and imaged using brightfield microscopy. The lesion cavity is clearly situated within the internal capsule. B, drawings of images from the left hemisphere, progressing from the most dorsal to most ventral to encompass the entire lesion. The image shown in (A) is highlighted. *AC = nucleus of anterior commissure; ACC = nucleus accumbens; ACX = decussation of anterior commissure; AQ = cerebral aqueduct; AS = arcuate sulcus; CA = caudate nucleus; CC = corpus callosum; CGS = cingulate sulcus; CTT = cortico-tectal tract; CS = central sulcus; GP = globus pallidus; GP i = globus pallidus internal division; GP e = globus pallidus external division; IC =internal capsule; IPS = intraparietal sulcus; Ins = insular cortex; LGd = dorsal lateral geniculate nucleus; LS = lateral sulcus; LV = lateral ventricle; Pu = putamen; Sb = subthalamic nucleus; SC = superior colliculus; TN = thalamic nuclei; 23 = cortical area 23; 3V = third ventricle*.

Figure 5B contains tracings from transverse sections, presented from most dorsal (top) to most ventral (bottom). Each section has been annotated with appropriate anatomical landmarks (Mikula et al., 2007) and contains the lesion cavity; depth increments relative to the previous section are also shown.

The lesion cavity extended approximately 17 mm along the dorso-ventral axis. In the most dorsal sections, the lesion was located within the caudate nucleus (CA) and formed a cavity of approximately 2 mm^2^ in area. In more ventral sections, the lesion increased in size (area approximately 6 mm^2^) and was primarily located within the internal capsule, with additional damage to the medial putamen (Pu) and lateral thalamic nuclei (TN). The internal capsule appeared diffuse in the more superior sections, but increasingly compact and V-shaped as it traveled ventrally through the basal ganglia. The lesion remained centered in the anterior half of internal capsule posterior limb (between the thalamus and globus pallidus), and extended laterally to the globus pallidus internus (GPi) and medially into the most lateral portion of the TN. In the most ventral sections, the lesion progressively decreased in size and ended at the level where the dorsal lateral geniculate nucleus (LGd) and subthalamic nucleus (Sb) were present.

These findings confirm that we successfully targeted the posterior limb subdivision of the internal capsule, thereby damaging a substantial portion of corticospinal fibers descending from various sensorimotor cortical regions. It is very likely the lesion also interrupted corticoreticular, corticorubral and corticopontine axons as these are also found in the anterior half of the posterior limb (Ara and Islam, 2010), as well as a small proportion of basal ganglia and thalamic circuits.

### Extent of RNm and Cortical Lesions

Histological analysis of these lesions has been presented in our previous report (Baines et al., 2026). In monkey Ca, two out of three electrolytic lesions were located within the RNm. Overall damage to layer V neurons across all cortical regions was the greatest in Monkey Ca (51%) compared to Monkey D (35%). Estimates of the damage to each cortical area are shown graphically in Fig. 6. Damage was widespread in both animals, encompassing PMd, both Anterior and Posterior divisions of Old M1, New M1, and S1 Area 3. In addition, the lesion in Monkey D extended into S1 Areas 1 and 2. Damage to Anterior Old M1, Posterior Old M1, New M1 and S1 Area 3 was approximately 2-fold greater in Monkey Ca than in Monkey D. In contrast, damage to PMd and S1 Areas 1 and 2 was considerable more in Monkey D. Monkey D did not have a preceding RNm lesion.

**Figure 6.**
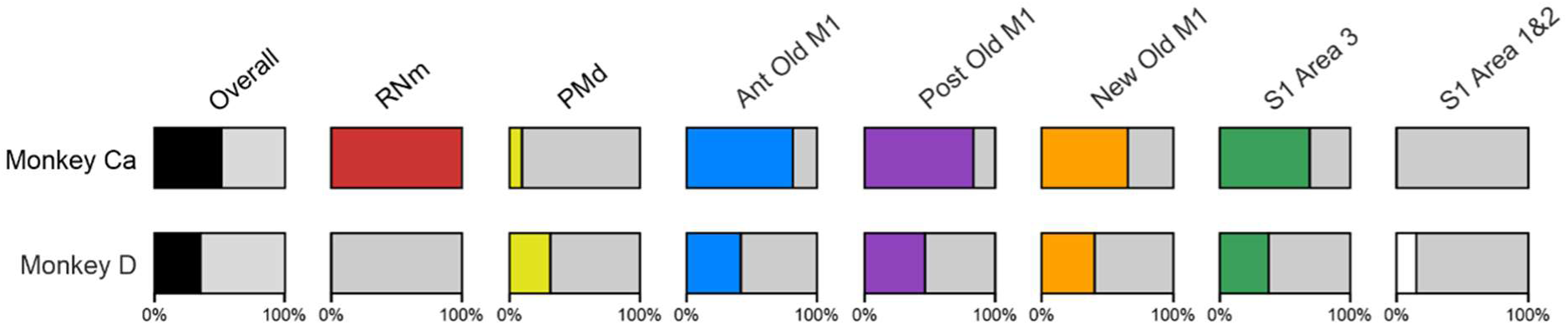
Estimated cortical lesion extent in Monkeys Ca and D. Bars show a visual representation of the percentage of layer V cells damaged in each cortical region. Overall values were calculated using an average of damage across all cortical regions in each monkey. RNm results have been expressed as 100% or 0% as there are no numerical values for this lesion type. Adapted from Baines et al. (2026).

### Impact of Lesions on Reaching Performance

Prior to receiving a lesion, the monkeys completed the task with ease, adopting an efficient and stereotyped path from handle to cup for both *in synergy* and *out of synergy* reaches (Fig. 7AB ‘baseline). Following a cortical lesion in Monkey D, the subacute phase demonstrated a slight impairment in task performance as the path from handle to cup became less direct (Fig. 7AB ‘subacute’). The paths shown correspond to the initial reach, detected as described in Methods, which was the focus of our analysis. The initial reach did not accurately reach the cup, requiring a subsequent sub-movement (not shown in Fig. 7). By the chronic period initial reach trajectories had largely returned to normal for the *out of synergy* reaches (Fig. 7A ‘chronic’), although trajectories remained more curved than at baseline for the *in synergy* reaches. In both cases the initial reach movements still fell short of the cup. By contrast, the internal capsule lesion in Monkey Cw resulted in significantly altered reaches, often characterized by repeated spiraling movements as the monkey progressed from handle to cup (Fig. 7CD). To quantify these changes in task performance and the underlying involvement of flexor synergies, we assessed flexion fraction strength, flexion fraction proportion and path length for the initial part of both *in synergy* and *out of synergy* reaches.

**Figure 7.**
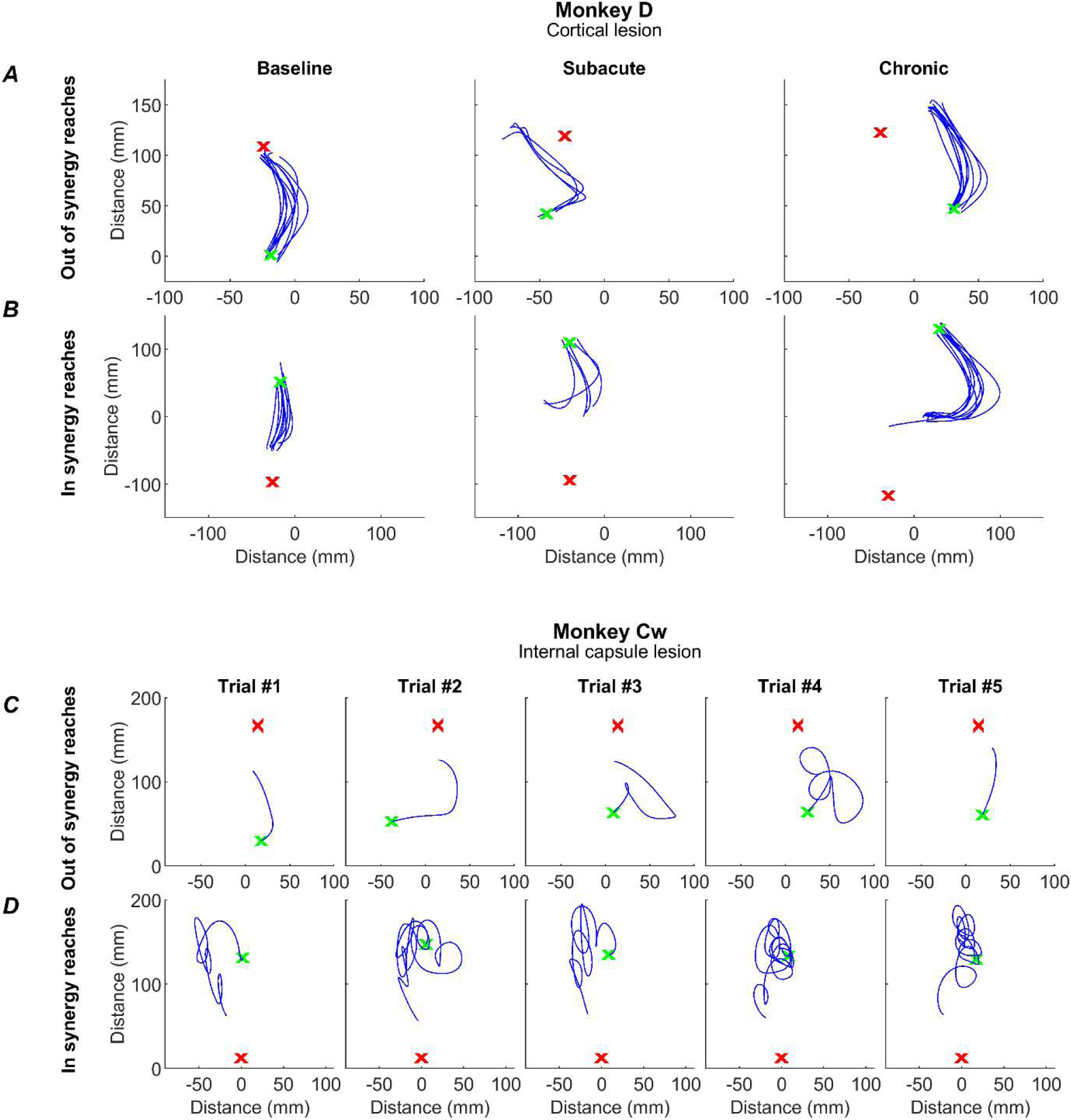
Example reaches. Reach trajectories, as determined by index finger tracking, for single trials (blue traces) for the initial reach (see Methods) from handle (green x) to cup (red x). For Monkey D, these are shown as overlays of all trials for an *out of synergy* (A) and *in synergy* (B) reach, for a baseline, subacute and chronic session (columns). Due to the increased variability in reach trajectory post-lesion in monkey Cw, each plot shows a single trial from a single post-lesion session in the Chronic period for an *out of synergy* reach (C) and an *in synergy* reach (D).

Figure 8 presents changes to three kinematic measures (rows) pre- and post-lesion for each monkey (columns). Results are presented for *out of synergy* (!Fig. 8ABC) and *in synergy* (Fig. 8DEF) movements. Flexion fraction proportion calculates how much of the movement is spent in flexor synergy (shoulder abduction with elbow flexion); flexion fraction strength calculates the strength of coupling between the shoulder and elbow joints during periods where the shoulder is abducting; path length provides a quantitative measure of reaching efficiency. Results are averaged across all trials performed in a given week, with post-lesion results compared to each animal’s baseline average.

**Figure 8.**
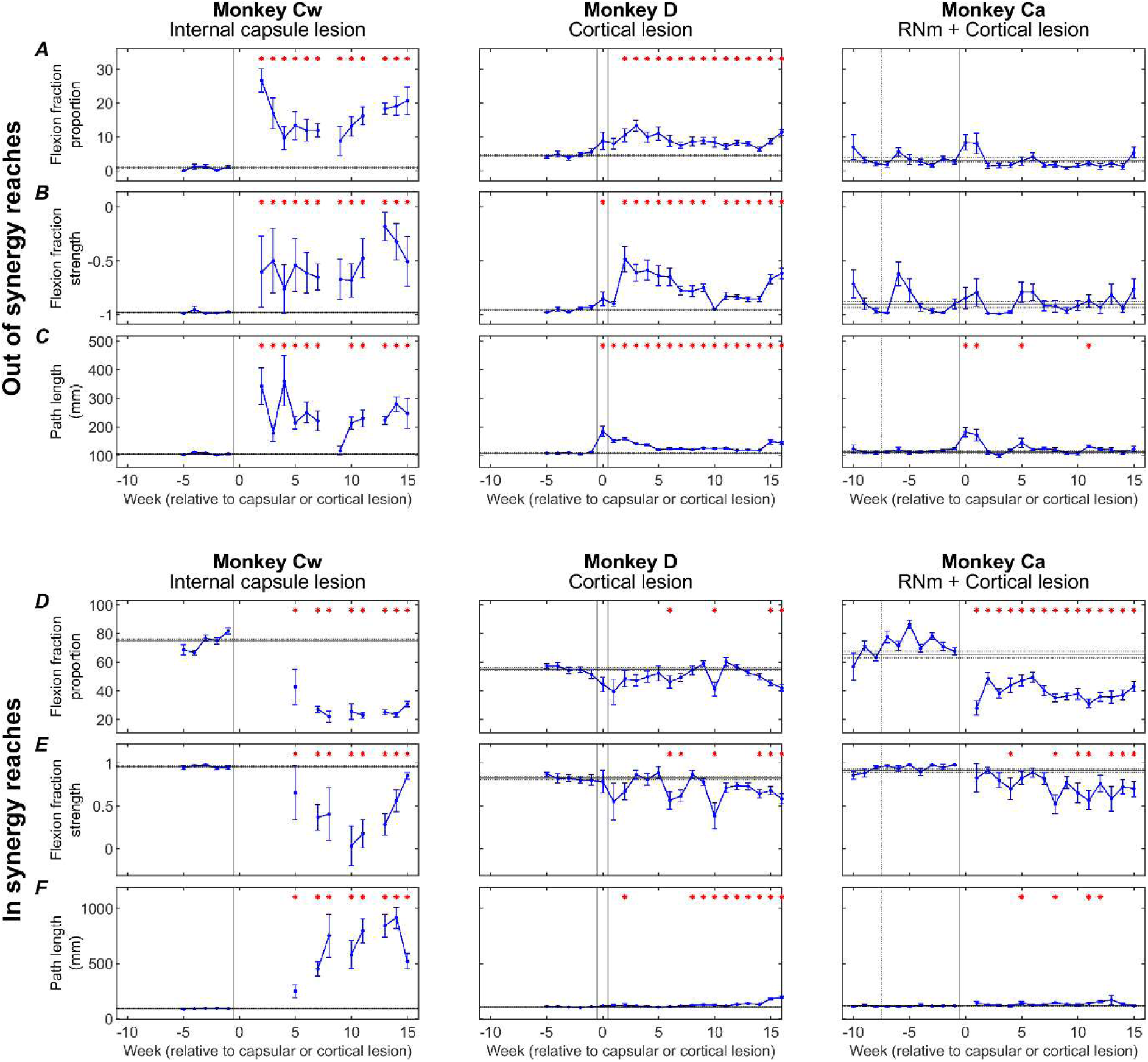
Weekly kinematic measures. Weekly flexion fraction proportion, flexion fraction strength, and path length for each monkey (columns), shown for *out of synergy* reaches (A-C) and *in synergy* reaches (D-F). Error bars are standard error, and red asterisks show a significant difference (p<0.05, adjusted by Benjamini-Hochberg procedure with a false discovery rate of 5%) relative to the pre-lesion period (horizontal lines show pre-lesion mean ± standard error). Dotted vertical lines show the RNm lesion date (Monkey Ca); vertical black lines show cortical (Monkeys Ca and D) or internal capsule lesion dates (Monkey Cw). Data gaps indicate insufficient trials for weekly analysis.

When examining *out of synergy* movements, Monkeys Cw and D showed significant increases in flexion fraction proportion (Fig. 8A) post-lesion, which showed limited recovery in both animals. During the chronic period (week 10 onwards), the proportion of time Monkey Cw spent in flexor synergy was approximately between two and three times larger than Monkey D. No significant increases in flexion fraction proportion were observed in Monkey Ca for *out of synergy* reaches.

A similar pattern of results was observed for flexion fraction strength (Fig. 8B). All monkeys began with a pre-lesion value of approximately -1 for healthy reaches, as expected for *out of synergy* reaches that required minimal elbow flexion coupled with shoulder abduction. This measure increased significantly only in Monkeys Cw and D. Flexion fraction strength was most abnormally altered in Monkey Cw, with mean R values ranging from -0.5 to 0 during the chronic period, compared to a less substantial change to between -0.8 and -0.5 for Monkey D over the same period. Again, this measure was not significantly changed in Monkey Ca post-lesion.

Path length showed immediate increases in all monkeys post-lesion (Fig. 8C). This eventually recovered in Monkey Ca, but remained elevated in Monkeys D and Cw. The greatest overall increases were seen in Monkey Cw.

When examining *in synergy* movements, all monkeys showed significant decreases in the flexion fraction proportion at some point during recovery (Fig. 8D). The largest decreases were observed in Monkey Cw, followed by Monkey Ca, with the smallest overall changes seen in Monkey D. Both Monkeys Cw and Ca showed limited, if any recovery of this measure.

Flexion fraction strength for *in synergy* movements (Fig. 8E) in each monkey’s baseline period was close to 1, as expected because these reaches require elbow flexion coupled with shoulder abduction. These values significantly decreased during lesion recovery in all monkeys, although the greatest reduction was observed in Monkey Cw, with R values ranging between 1 to 0, compared with ranges of 1 to 0.5 for Monkeys Ca and D.

Path length was significantly increased following the lesion in all monkeys (Fig. 8F); however, the magnitude of change was considerably greater in Monkey Cw than in Monkeys D and Ca. This eventually recovered in Monkey Ca, but not for Monkeys Cw and D.

Following the RNm lesion alone in Monkey Ca (period between the dotted and solid vertical lines), none of the measures were significantly changed during either *out of synergy* or *in synergy* reaches.

Figure 9 presents the same three kinematic assessments averaged across pre-lesion, acute, and chronic recovery periods to highlight the overall significant trends.

**Figure 9.**
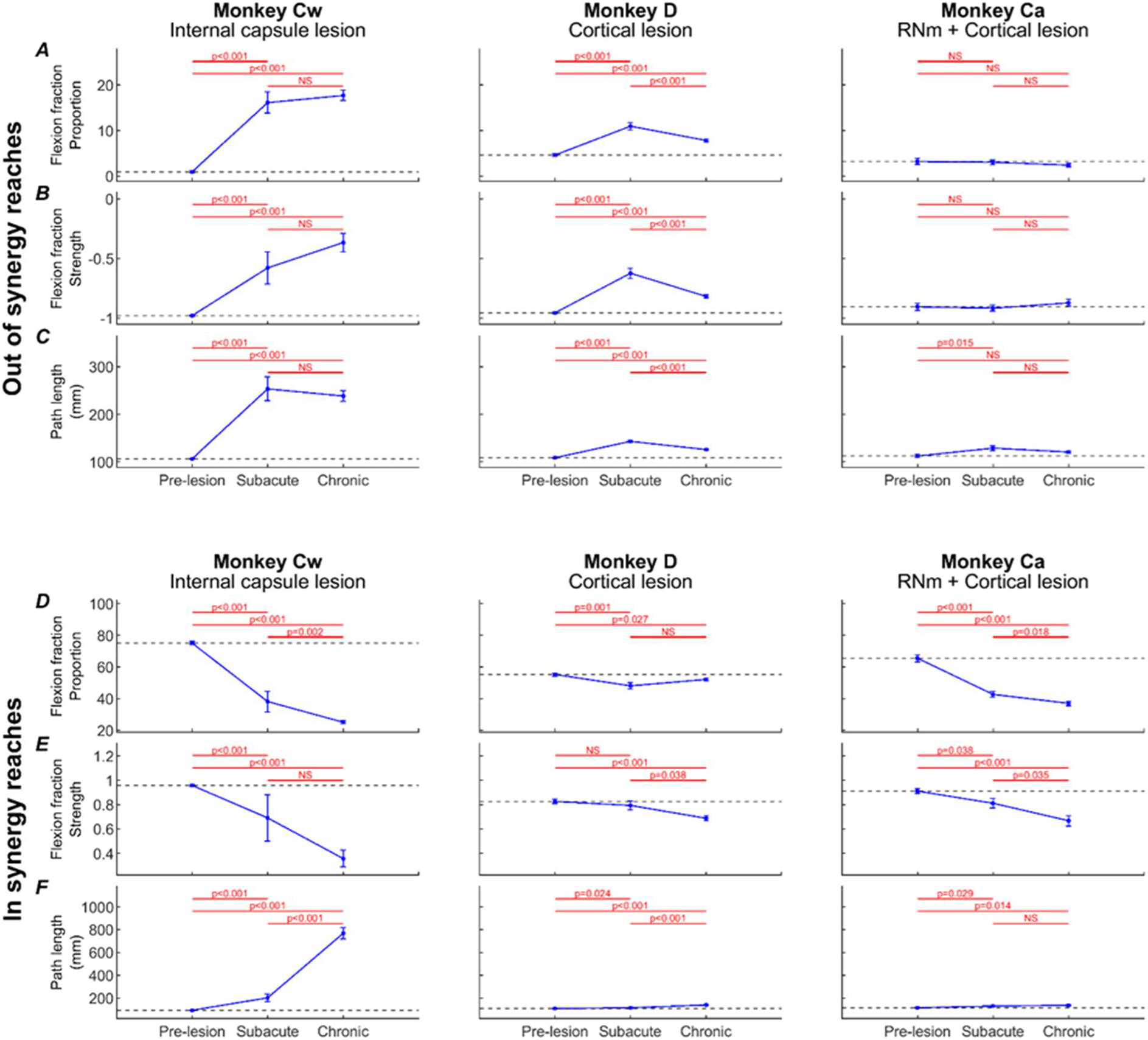
Pre-lesion, subacute and chronic kinematic measures. Flexion fraction proportion (A,D), flexion fraction strength (B,E), and path length (C,F) for each monkey (columns), shown for *out of synergy* reaches (A-C) and *in synergy* reaches (D-F), averaged across pre-lesion, subacute (0-5 weeks) and chronic (10-15 weeks) periods. Error bars are standard error. Significance testing was performed between pairs of value using unpaired t-tests, with p-values adjusted by Benjamini-Hochberg procedure with a false discovery rate of 5% (NS: non-significant, p>0.05). Black dotted lines show pre-lesion means.

During *out of synergy* movements (Fig. 9ABC), Monkey Cw showed significant acute increases in flexion fraction proportion (Fig. 9A), flexion fraction strength (Fig. 9B), and path length (Fig. 9C) following the internal capsule lesion; these increases persisted into the chronic period with no significant recovery. Monkey D had significant acute increases in all metrics, which did show significant recovery by the chronic period (unlike Monkey Cw), although all three measures remained significantly elevated relative to baseline. By contrast, Monkey Ca did not show any significant increases in flexion fraction proportion or strength during *out of synergy* movements, with the acutely increased path length returning to baseline by the chronic period. Together, these findings indicate that intrusion of synergies occurred in Monkeys Cw and D, most prominently in Monkey Cw, whereas Monkey Ca did not exhibit this effect.

By comparison, *in synergy* movements (Fig. 9DEF) were affected across all monkeys, shown by significantly reduced flexion fraction proportion (Fig. 9D) and flexion fraction strength (Fig. 9E), and significantly increased path length (Fig. 9F) by the chronic periods. None showed signs of recovery from the acute to chronic periods for any measure. Given this universal outcome with *in synergy* reaches, these results likely reflect a general motor deficit rather than the synergy-specific effects observed in *out of synergy* reaches.

To dissect the directionality of impairments further, we quantified trial initiation attempts (Fig. 10). Results are shown for each monkey (columns), comparing trial types (rows) that required elbow flexion to hold the handle (‘handle near’ – Fig. 10A), versus trials requiring elbow extension (‘handle far’ – Fig. 10B); these results are averaged by week. Figure 10CD presents the same data and arrangement (‘handle near’ – Fig. 10C; ‘handle far’ – Fig. 10D), but now averaged over all trials within the baseline, subacute, and chronic recovery periods. If flexor synergy were expressed, we would expect a higher number of handle attempts during ‘handle far’ trials, as these would require overcoming the flexor synergy to extend the elbow while abducting the shoulder in order to make contact with the handle.

**Figure 10.**
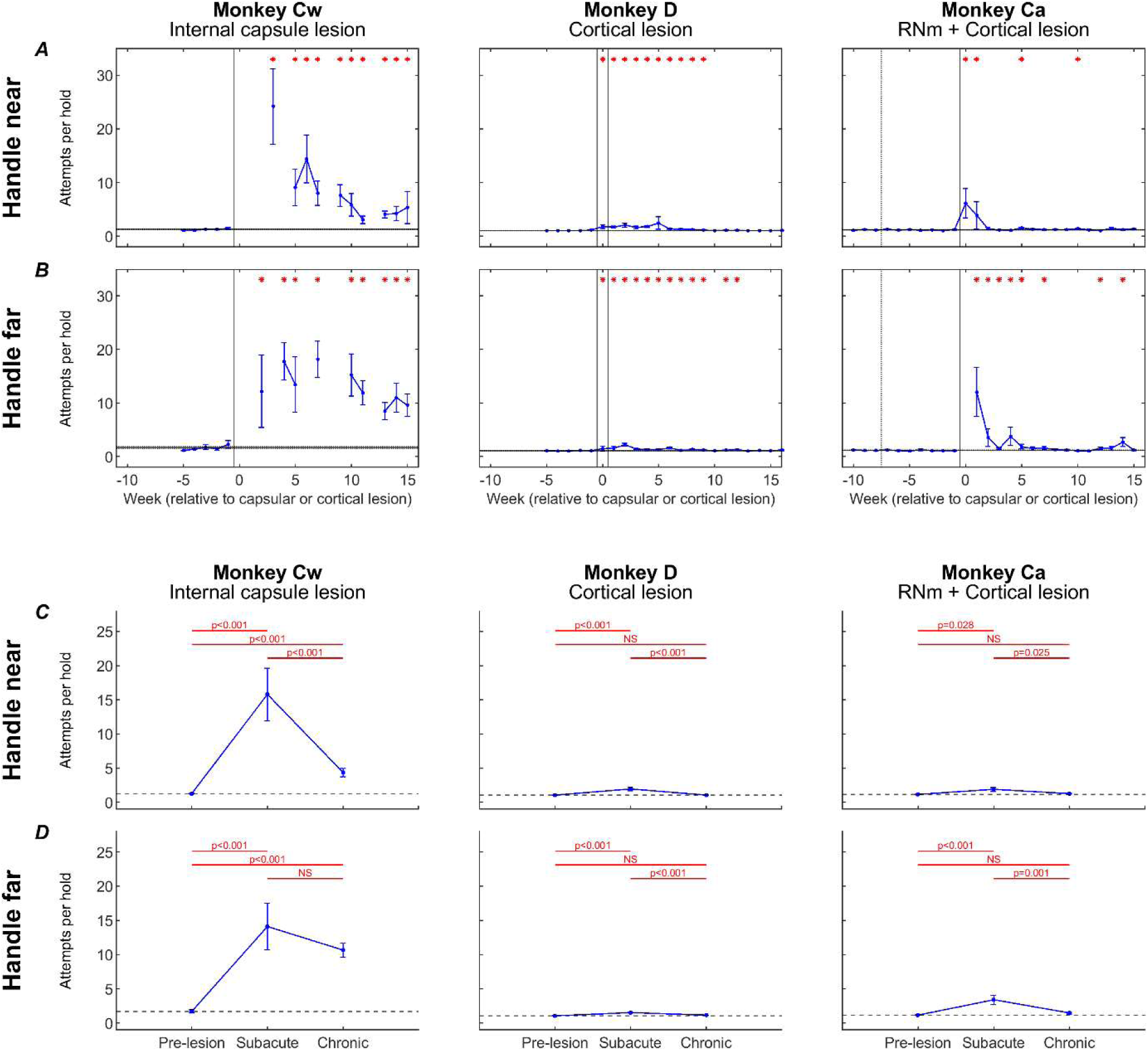
Trial initiation attempts. Number of attempts required to initiate a reaching trial (see Methods), for trials with the handle positioned near the body (*in synergy*; A,C) or far from the body (*out of synergy*; B,D) shown as weekly averages (A,B) and averages across pre-lesion, subacute and chronic (C,D) periods. Error bars show standard error. A,B, red asterisks show a significant difference (p<0.05, adjusted by Benjamini-Hochberg procedure with a false discovery rate of 5%) relative to the pre-lesion period (horizontal lines show pre-lesion mean ± standard error). Dotted vertical lines show the RNm lesion date (Monkey Ca); vertical black lines show cortical (Monkeys Ca and D) or internal capsule lesion dates (Monkey Cw). Data gaps indicate insufficient trials for weekly analysis. C,D, significance testing was performed between pairs of values using unpaired t-tests, with p-values adjusted by Benjamini-Hochberg procedure with a false discovery rate of 5% (NS: non-significant, p>0.05). Black dotted line shows pre-lesion means.

Before each lesion, all monkeys were able to initiate each trial with very few (typically just one) handle hold attempts. Following the internal capsule lesion in Monkey Cw, there was a large and significant increase in the number of attempts to grasp the handle for both ‘handle near’ (*in synergy* – Fig. 10AC) and ‘handle far’ (*out of synergy* – Fig. 10BD) trials, with an average of 15 attempts per trial during the subacute period. By the chronic period, ‘handle near’ trials became significantly easier to initiate, requiring an average of five attempts per trial; by contrast, ‘handle far’ trials did not show significant improvement, remaining at 11 attempts per trial on average. This pattern provided strong evidence for the expression of flexor synergy in Monkey Cw, which showed poor recovery.

In Monkeys D and Ca, smaller, but still significant, increases in handle-hold attempts were observed during the subacute recovery period for both ‘handle near’ (Fig. 10C) and ‘handle-far’ trials (Fig. 10D). By the chronic period, performance of both trial types had significantly recovered in both animals. These results represent the less severe recovery outcomes seen in these animals compared to Monkey Cw.

### Quantifying Flexor Synergy Expression with EMG Cross Correlation

In addition to quantifying abnormal kinematics, we also performed cross correlation analyses of EMG signals recorded from different muscle groups during reaching movements (Fig. 11). The coefficient of determination (r²) was used as a measure of co-contraction for each muscle pair. For *out of synergy* reaches (left column), the average r^2^ per week of recordings was calculated using EMG from available shoulder abductor muscles (anterior deltoid, posterior deltoid, supraspinatus) and elbow flexors (biceps, brachioradialis and brachialis). For *in synergy* reaches (right column), the same method was applied using EMG from an available shoulder abductor (posterior deltoid) and an elbow extensor (triceps). Thus, the muscle synergies analyzed for each reach type correspond to the opposite of those selectively deployed during healthy movements; in essence, any increases in r^2^ relative to baseline indicate abnormal synergies.

**Figure 11.**
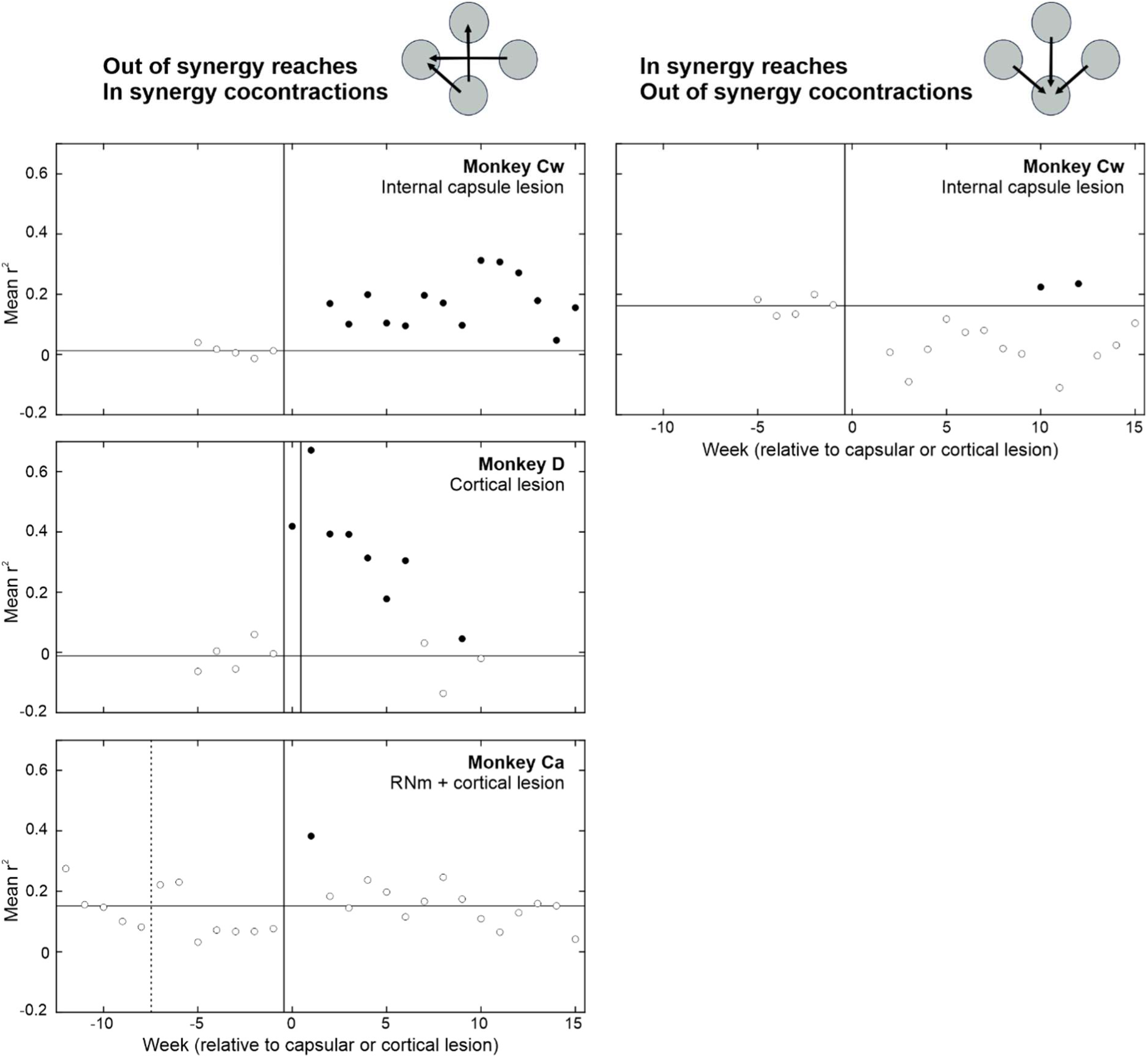
Quantification of abnormal muscle co-contractions during *out of synergy* vs *in synergy* reaching using EMG cross correlation. Rows show results from different monkeys. Muscle synergies analyzed for each reach type correspond to the opposite of those typically deployed during healthy movements performance. For *out of synergy* reaches (left), cross correlation coefficients were calculated between EMG activity from available shoulder abductor muscles (anterior deltoid, posterior deltoid and supraspinatus) and elbow flexors (biceps, brachioradialis and brachialis). For *in synergy* reaches (right), the same method was applied using EMG from an available shoulder abductor (posterior deltoid) and an elbow extensor (triceps). Dotted vertical lines show the RNm lesion date (Monkey Ca); vertical black lines show cortical (Monkeys Ca and D) or internal capsule lesion dates (Monkey Cw). Horizontal lines indicate baseline average coefficient of determination (r^2^). Y axis ranges are the same across all plots to allow for easier visual comparison. Filled circles are significantly different from baseline (p < 0.05), identified using the two-sample t test. *In synergy* analyses were not possible in Monkeys D and Ca due to technical issues with triceps EMG recordings. EMG recording was not possible past week 10 in Monkey D due to technical issues with the headpiece implant.

Following the internal capsule lesion in Monkey Cw, there were significant increases in r^2^ values for *in synergy* co-contractions (i.e. flexor synergy) when performing *out of synergy* reaches that normally require elbow extension. This flexor synergy persisted into the chronic recovery period. In contrast, no corresponding increase was observed for *out of synergy* co-contractions during *in synergy* reaches (i.e. the opposite of flexor synergy), indicating a selective expression of the flexor synergy in this animal which did not recover.

Following the first cortical lesion in Monkey D, there were large, significant increases in r^2^ values for *in synergy* co-contractions during *out of synergy* reaching; this effect was further exacerbated by the second lesion. Increased r^2^ values were transient and returned to baseline levels by the chronic recovery period (week 10). Analyses of *in synergy* muscle co-contractions was not possible due to technical issues with the triceps EMG recording. As a result, we cannot conclude from these data alone that the observed co-contractions reflected obligate flexor synergy expression. Nevertheless, consistent with the kinematic findings, Monkey D showed more successful recovery than Monkey Cw.

Following the RNm lesion in Monkey Ca, no significant changes to *in synergy* co-contraction strength were observed during *out of synergy* reaches. The subsequent cortical lesion caused a transient increase in r^2^ which resolved by week 2 of recovery. Analysis of *in synergy* muscle co-contractions was not possible in Monkey Ca due to technical issues with triceps recording. Overall, these results indicate Monkey Ca did not exhibit the same degree of impairment attributable to positive signs as observed in Monkeys Cw and D.

There was an important congruence between the changes seen in muscle co-contraction during *out of synergy* reaches in Fig. 11 and those observed in the kinematic measures of Fig. 8AB. On both measures, Monkey Cw showed a persistent increase, Monkey D a transient increase which recovered, and Monkey Ca a very brief rise which had normalized after two weeks. This congruence between metrics derived from very different underlying signals provides confidence that they are both measuring the same underlying phenomenon of the flexor synergy.

## Discussion

### Relative Differences in Abnormal Flexor Synergy Expression following Internal Capsule vs Focal Cortical Lesions

In this study, we show that a large, unilateral lesion to the internal capsule generated severe abnormal flexor synergy with little to no meaningful recovery into the chronic phase (Monkey Cw). In contrast, a widespread cortical lesion involving multiple sensorimotor regions (PMd, M1, S1) also caused abnormal flexor synergy, but the impairment was comparatively less severe and showed significant recovery by the chronic phase (Monkey D). Flexor synergy did not emerge following combined lesions to the left RNm and multiple sensorimotor cortical regions (PMd, M1, S1; Monkey Ca). A summary of these findings is presented in Table 1.

**Table 1.** Summary of different lesion effects.

| Lesion | Flexion fraction proportion and strength | Trial initiation attempts | *In synergy* co-contractions | *Out of synergy* co-contractions | Flexor synergy |
| --- | --- | --- | --- | --- | --- |
| **RNm + Cortical Lesion (Monkey Ca)** | Changes observed for *in synergy* reaches which never recovered.<br>*Out of synergy* reaches unaffected | Near-normal initiation of trials which required *out of synergy* ('handle far') and *in synergy* ('handle near') movement | Small transient increase immediately post-lesion which recovered within 2 weeks | - | None |
| **Cortical Lesion (Monkey D)** | Changes observed for *in synergy* and *out of synergy* reaches.<br>*Out of synergy* reaches showed much poorer recovery | Near-normal initiation of trials which required *out of synergy* ('handle far') and *in synergy* ('handle near') movement | Large transient increases which recovered within 9 weeks | - | Mild expression |
| **Internal Capsule (Monkey Cw)** | Largest changes to *in synergy* and *out of synergy* reaches in study.<br>Both showed very poor recovery | Very poor initiation of trials which required *out of synergy* movement ('handle far') compared to *in synergy* movement ('handle near') | Large, persistent increases with very poor recovery | No consistent increases | Severe expression |

This work demonstrates the utility of sensitive, three dimensional kinematic measures to quantify synergy intrusion during a naturalistic reaching task similar to that previously described by Avni and colleagues (Avni et al., 2024a). In an earlier study (Baines et al., 2026), we noted that Monkey D did not express overt synergies (detectable by clinical observation), emphasizing the importance of sensitive quantification techniques that enable accurate and relative comparisons of synergy expression.

The internal capsule lesion in Monkey Cw produced severe abnormal synergies, evidenced by kinematic and EMG changes consistent with the characteristic flexor synergy pattern, which showed no signs of recovery. This result can be explained considering the affected anatomy in Monkey Cw. Lesion damage involved the posterior limb of the internal capsule; as intended, few, if any, corticospinal axons would have been spared. In addition to corticospinal fibers, the posterior limb also contains corticoreticular, corticorubral and corticopontine projections (Kuypers, 1960; Emos et al., 2023) which were also likely disrupted. The lesion appeared to extend partially into the genu and anterior limb of the internal capsule more rostrally, likely resulting in damage to corticofugal outflow from the supplementary motor area (SMA) and premotor regions respectively (Fries et al., 1993; Shelton and Reding, 2001).

The lesion in Monkey Cw also generated partial but significant damage to adjacent structures, including components of the basal ganglia (globus pallidus, putamen, and caudate nucleus), as well as more anterior thalamic nuclei. Clinically internal capsule strokes often extend into these neighboring structures (Miyai et al., 1997; Shelton and Reding, 2001). Involvement of the basal ganglia either in isolation or in association with capsular strokes does not appear to influence upper limb recovery outcomes in humans, whereas involvement of the thalamus in combination with damage to the posterior limb of the internal capsule lesions has been shown to impact recovery negatively (Fries et al., 1993; Shelton and Reding, 2001). In Monkey Cw, a large proportion of thalamic structures remained intact following the lesion. Therefore, the most likely explanation for the severe and persistent motor deficits observed is the disruption of corticofugal pathways. However, a potential contribution of thalamic and basal ganglia damage cannot be entirely excluded.

It is unlikely that any viable corticofugal routes remained to support recovery from the ipsilesional cortex following the internal capsule lesion. In this situation, only three surviving pathways could subsume control of the affected upper limb; all arise from the contralesional cortex: 1) corticoreticular connections and the bilateral reticulospinal tract (Riddle et al., 2009; Zaaimi et al., 2012), 2) the ipsilateral corticospinal tract from contralesional cortex (Soteropoulos et al., 2011; Morecraft et al., 2016) and 3) the contralateral corticospinal tract from contralesional cortex, connecting to the affected side motoneurons via commissural interneurons (Jankowska et al., 2006; Maxwell and Soteropoulos, 2020). Importantly, all of these pathways rely predominantly on oligosynaptic activation of motoneurons. Their contribution to voluntary motor output would presumably be much less if the ipsilesional corticospinal tract were intact. Several previous studies have implied that greater involvement of the contralesional cortex is correlated with poorer upper limb recovery in human stroke patients (Turton et al., 1996; Ward et al., 2003; Matsuura et al., 2017; McPherson et al., 2018a; Karbasforoushan et al., 2019).

Previous experiments using microstimulation within the intermediate zone of the spinal cord in healthy macaques revealed that only 7% of sites were able to activate shoulder abductor muscles without elbow flexors. The majority of cervical spinal cord interneurons seem capable only of facilitating *in synergy* co-contractions (Glover et al., 2025). In contrast, microstimulation of contralateral M1 and the reticular formation was able to recruit shoulder abductor and elbow flexor muscles selectively. This study also showed that neurons in motor cortical areas and the reticular formation showed a preference to fire in co-contractions orthogonal to post-stroke synergies (e.g. shoulder abduction/elbow extension or shoulder adduction/elbow flexion) (Glover et al., 2025). These data argue against the notion that supraspinal areas are themselves the primary generator of flexor synergy following severe cortical damage, contrary to previous hypotheses in the field (Ward et al., 2006; Owen et al., 2017; McPherson et al., 2018a).

It is plausible that, following the internal capsule lesion, motor commands are forced to be routed through intact motor pathways originating from the contralesional cortex. In contrast to the selective monosynaptic cortico-motoneuronal connections destroyed by the lesion, these surviving pathways must access motoneurons via spinal cord interneuron circuits that preferentially facilitate *in synergy* co-contractions regardless of the motor task. This maladaptive recruitment of indirect motor pathways likely represents the only feasible means of generating any upper limb movement, albeit with long-lasting loss of isolated joint coordination in Monkey Cw.

After a partial cortical lesion (Monkey D), the flexor synergy was less severe in the acute phase, and demonstrated substantially greater recovery compared to Monkey Cw. It is plausible that the acute synergy expression in this case was mediated by similar maladaptive pathways, however, over time, these patterns may have come under increasing control of the surviving ipsilesional cortex. Notably, the lesion in Monkey D did not exceed more than 50% damage to ipsilesional PMd, M1, or S1, and SMA remained entirely intact. SMA preservation is often seen in clinical practice following middle cerebral artery strokes, as it is supplied by the anterior cerebral artery (Miyai et al., 1999). Monosynaptic connections with motoneurons from the spared ipsilesional cortex (particularly New M1) (Rathelot and Strick, 2009; McNeal et al., 2010; Witham et al., 2016), SMA (Jürgens, 1984; Liu and Rouiller, 1999; McNeal et al., 2010), the reticular formation (Riddle et al., 2009; Boubker Zaaimi, 2012), and the RNm (Ralston et al., 1988; Miller et al., 1993; Belhaj-Saïf et al., 1998) could have retained sufficient plastic potential to reestablish fractionated control over muscles in this case.

Preservation/restoration of selective ipsilesional drive could explain why chronic phase improvements were observed in Monkey D, but not in Monkey Cw. This interpretation is consistent with findings from Shelton and Reding (2001), who reported that stroke patients were substantially more likely to recover at least some movement following purely cortical strokes compared to subcortical or mixed strokes. Of those that recovered some movement, damage to the PLIC was highly predictive of whether patients remained either in synergy, or were capable of making isolated joint movements (20% vs 83% isolated movements for PLIC damaged vs spared respectively). Our findings are also consistent with studies demonstrating patients with strokes confined to the internal capsule and basal ganglia exhibit a diminished response to rehabilitation compared to those with damage isolated to the cortex (Pantano et al., 1995; Miyai et al., 1997). It appears that Monkey D followed a more classical trajectory of motor recovery, consistent with the staged progression described by Twitchell and Brunnstrom (Twitchell, 1951; Brunnstrom, 1966, 1970), whereas Monkey Cw resembles patients who plateau at the stage of obligate synergy expression.

### No Abnormal Flexor Synergy Expression following a Combined RNm and Cortical Lesion

In our previous study (Baines et al., 2026), we showed that combined lesions to the RNm and the sensorimotor cortex in Monkey Ca led to worse recovery of upper limb reaching speed relative to monkeys with lesions confined to the cortex. These findings reaffirmed the significant contributions of the rubrospinal system in supporting voluntary movement and recovery in monkeys (Cheney, 1980; Holstege et al., 1988; A Belhaj-Saïf, 1998; Belhaj-Saïf and Cheney, 2000). In contrast, the rubrospinal tract is considered comparatively vestigial in humans (Hatschek, 1907; Nathan and Smith, 1955; Donkelaar, 1988; Massion, 1988; Basile et al., 2021). Here we hypothesized that the rubrospinal tract originating in the RNm could provide monkeys with an additional route for recovery from synergies, potentially explaining why monkeys typically recover from lesions better than humans.

In this study, combined lesions to the RNm and sensorimotor cortex did not produce flexor synergy expression. Rather, the opposite effect was observed: significant and persistent changes in kinematic measures were evident only during *in synergy* reaching. No increase of *in synergy* co-contractions (shoulder abductors with elbow flexors) during *out of synergy* reaching was seen following the cortical lesion (unlike Monkeys D and Cw). This finding is consistent with our previous analysis in monkey Ca, in which movements requiring elbow extension (*out of synergy*) showed better recovery of reaching speed and trajectory variability than those requiring elbow flexion (*in synergy*) (Baines et al., 2026). The opposite phenotype is typically seen in human stroke survivors, with biased recovery of flexors and persistently weakened extensors (Kamper et al., 2003; Baker et al., 2015). This is a particularly interesting result given that the same cortical regions that affected Monkey D (PMd, M1, and S1) were also involved in monkey Ca, with an even greater extent of layer V damage across M1. Despite the more extensive loss of corticofugal neurons in Monkey Ca, abnormal flexor synergy did not emerge. Although we have previously shown that RNm lesions impair recovery of trajectory variability and speed after cortical lesions, it seems that the rubrospinal tract is not especially important in preventing synergies in macaques.

One possible explanation relates to the functional properties of the RNm in health versus following corticospinal loss. In healthy animals, rubrospinal outputs preferentially facilitate extensor muscles (Mewes and Cheney, 1991; Belhaj-Saïf et al., 1998); however, following unilateral pyramidotomy, these outputs shift towards cofacilitation of both flexor and extensor motoneurons (Belhaj-Saïf and Cheney, 2000). This reorganization likely reflects compensatory recruitment of the rubrospinal system to restore lost corticospinal drive, particularly to flexor muscles. In Monkey Ca, lesioning the RNm would have prevented such compensatory adaptation, potentially causing imbalanced facilitation of extensor muscles and poorer ability to perform *in synergy* reaches as a result. This could also explain the different recovery outcomes in Monkey Ca compared to Monkey D.

It is important to note that the lesion in Monkey Ca, like Monkey D, spared a portion of corticofugal outputs from M1, PMd, and S1, and left SMA entirely intact. These surviving pathways likely provided sufficient ipsilesional control over synergistic spinal circuits in this animal, preventing the expression of the flexor synergy.

All three animals also showed impairments to *in synergy* reaches, with significant loss of the normal shoulder-elbow coordinated flexion seen at baseline, and a significant increase in path length (Fig. 8DEF, 9DEF). Importantly, these changes did not seem to relate simply to those during *out of synergy* reaches. They were largest in Monkey Cw, which also had the largest *out of synergy* changes corresponding to the flexor synergy. However, *in synergy* flexion fraction abnormalities were greater in Monkey Ca than Monkey D, whereas for *out of synergy* reaches the converse was true. We have previously shown that Monkey Ca showed larger and more persistent deficits on trajectory variability and maximum speed than Monkey D (Baines et al., 2026), reflecting greater disruption to descending control. The widespread changes to *in synergy* probably reflect similar generalized impairment. On this view, it is unsurprising that Monkey Cw should show the greatest effects to *in synergy* as well as *out of synergy* reaches, as the damage to the corticospinal tract in this animal was greater than in the other two.

### Conclusion

This study reveals clear differences in recovery trajectories following three distinct lesion types: a focal cortical lesion, a combined cortical and RNm lesion, and a lesion to the internal capsule. A large lesion confined to the sensorimotor cortex produced mild flexor synergy detected only through the use of sensitive quantitative analyses. Because this lesion spared portions of the upper limb representation within affected cortical areas, and left SMA entirely intact, the surviving ipsilesional corticofugal pathways were likely able to strengthen and sculpt their monosynaptic connections to synergistic spinal circuits over time, hence facilitating recovery. In contrast, combining an RNm lesion with a cortical lesion (involving the same sensorimotor regions) did not result in flexor synergy expression. A large lesion to the internal capsule produced overt flexor synergy that showed little recovery. The internal capsule lesion likely eliminated the majority of corticofugal connectivity from the ipsilesional cortex, leaving spinal circuits only capable of *in synergy* activation as the only viable means to drive movement on the affected side. Collectively, these findings indicate that the emergence of abnormal synergies is determined by the extent of corticofugal disruption, and the capacity of surviving supraspinal motor pathways to regain selective control over biased spinal circuits.

## Acknowledgements

The authors wish to thank Andrew Atkinson, Terri Jackson and Stevie O’Keefe for animal training; Norman Charlton for engineering support; Kathy Murphy, Rocio Palacios O’Connor, Fiona Douglas, Ines Sanchez Garcia, and Daniel Blake for veterinary assistance; and Michelle Waddle for surgical theatre nursing.

## Funding

This work was supported by NIH grant 5R01NS119319.

## Notes

### Competing Interest Statement

The authors have declared no competing interest.

### Summary of Updates

Citation of Baines et al (2026) corrected in the reference list.

